# QuantifyPolarity, a new tool-kit for measuring planar polarized protein distributions and cell properties in developing tissues

**DOI:** 10.1101/2020.11.24.396044

**Authors:** Su Ee Tan, Weijie Tan, Katherine H. Fisher, David Strutt

## Abstract

The coordination of cells or structures within the plane of a tissue is known as planar polarization. It is often governed by the asymmetric distribution of planar polarity proteins within cells. A number of quantitative methods have been developed to provide a readout of planar polarization of protein distributions. However, the quantification of planar polarization can be affected by different cell geometries. Hence, we developed a novel planar polarity quantification method based on Principal Component Analysis (PCA) that is shape insensitive. Here, we compare this method with other state-of-the-art methods on simulated models and biological datasets. We found that the PCA method performs robustly in quantifying planar polarity independently of variation in cell geometry. Furthermore, we designed a user-friendly graphical user interface called QuantifyPolarity, equipped with three different methods for automated quantification of polarity. QuantifyPolarity also provides image analysis tools to quantify cell morphology and packing geometry, allowing the relationship of these characteristics to planar polarization to be investigated. This all-in-one tool enables experimentalists with no prior computational expertise to perform high-throughput cell polarity and shape analysis automatically and efficiently.

## Introduction

Planar polarity plays a crucial role in coordinating the behavior of groups of cells to generate highly organized structures at the level of tissues and organs. It governs oriented cell divisions and cell rearrangements that specify tissue shape and generates global alignment of external structures such as *Drosophila* wing hairs, mammalian hair and cilia, bristles covering the insect epidermis or reptilian scales covering the body surface, and stereocilia bundles in the organ of Corti (Butler and Wallingford, 2017; Devenport, 2016).

Coordination of cell behavior at the tissue level is often initiated by polarization of proteins at the molecular level. Examples of planar polarization are: (1) Asymmetric cellular localization of core planar polarity proteins such as Frizzled (Fz), which determine the placement of *Drosophila* pupal wing hairs at distal cell edges (Strutt, 2001) (Figure 1A-C). (2) The Fat-Dachsous (Ft-Ds) system, in which intracellular asymmetry of Ft-Ds heterodimers (Brittle et al., 2012; Ambegaonkar et al., 2012; Bosveld et al., 2012; Merkel et al., 2014) results in the asymmetric distribution of the atypical myosin Dachs (Brittle et al., 2012; Mao et al., 2006), which has been implicated in directing the orientation of cell divisions in *Drosophila* wing discs (Mao et al., 2011) and cell rearrangements in the notum (Bosveld et al., 2012). (3) Planar polarization of proteins such as non-muscle Myosin II, filamentous actin (F-actin), and myosin activators Rho kinase (Rok), RhoA GTPase and Shroom, and cell adhesion proteins and regulators such as E-Cadherin, Bazooka (Par-3) and Armadillo (β-catenin) which are required for active cell contractility and rearrangements during *Drosophila* germ-band extension (Zallen and Wieschaus, 2004; Bertet et al., 2004; Blankenship et al., 2006; Simões et al., 2010; Simões et al., 2014; Kasza et al., 2014; Tetley et al., 2016; Tamada et al., 2012; Levayer and Lecuit, 2013).

**Figure 1.**
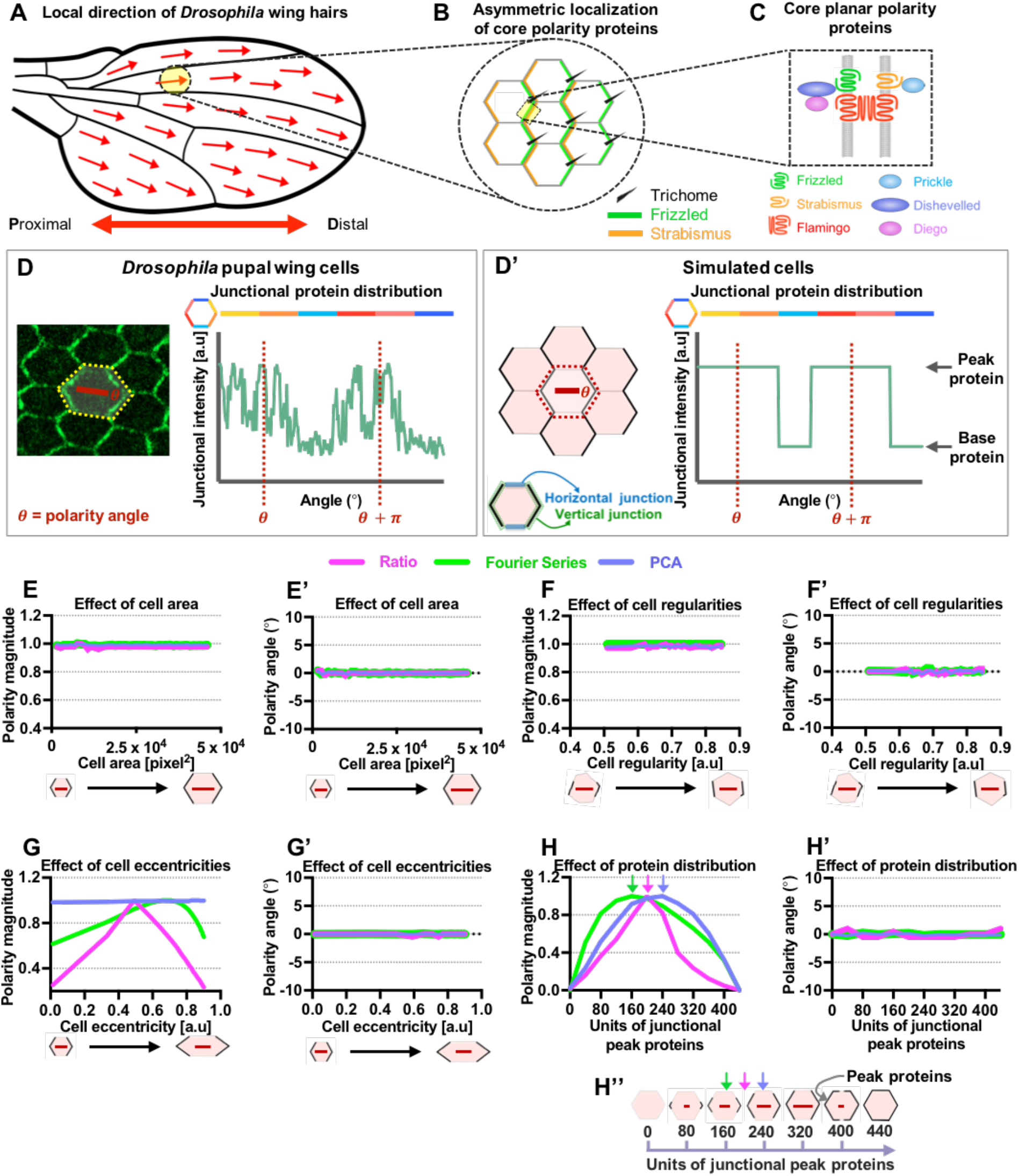
The PCA method is insensitive to variation in simulated cell sizes, regularities and eccentricities. (A) Cartoon of *Drosophila* wing blade. The red arrows indicate the local hair orientation. (B) Core planar polarity proteins asymmetrically localize on the opposite sides of the cell, with trichomes (hairs) forming from the distal edge colocalizing with Frizzled (green). Only the localizations of Frizzled and Strabismus (orange) on the cell junctions are shown. (C) Asymmetric distribution of core planar polarity pathway components along the proximodistal axis at the apical cell boundaries. (D-D’) Examples of junctional protein distribution of polarized (D) pupal wing cell and (D’) simulated cell with horizontal junctions (represented by blue box) and vertical junctions (represented by green box). For all simulated cells, peak protein is defined as proteins with higher intensity (255 a.u.) while base protein referred to proteins with lower intensity (40 a.u.). (E-H) Comparison between three methods of planar polarity quantification (Ratio, Fourier Series and PCA) using simulated cells with varying properties. (E-E’) Quantified polarity magnitudes (E) and polarity angles (E’) of cells with varying apical area, from 1500 pixel^2^ to 46000 pixel^2^ (~10 to 300 μm^2^). (F-F’) Quantified polarity magnitudes (F) and polarity angles (F’) of cells with varying shape regularity, from 0.5 to 0.85. (G-G’) Quantified polarity magnitudes (G) and polarity angles (G’) of cells with varying eccentricity, from 0 to 0.9. (H-H’’) Graphs show quantified polarity magnitudes (H) and polarity angles (H’) of cells with varying junctional protein distribution. Given a cell with total perimeter of 440 pixels, units of junctional peak proteins increase gradually (from 80 to 440 pixels) starting from both poles of vertical junctions. In Figure (H) and (H’’), arrows are pointing at unit of junctional peak proteins which gives maximum polarity magnitude for each method (magenta – Ratio, green – Fourier Series and blue – PCA). In this figure, all polarity magnitudes (a.u.) obtained using each different method are normalized. All polarity angles (in degree) range between −90° to +90°, with 0° corresponding to the x-axis of the image.

Planar polarity in epithelia varies as cell morphology changes dynamically throughout development (Classen et al., 2005; Aigouy et al., 2010), but how cells with different shapes and sizes achieve planar polarization is an important question that is difficult to address. This is in part due to the lack of a robust quantification method to quantify planar polarity independently of cell geometry. Currently available methods of planar polarization quantification suffer some limitations. Generally, existing methods can be classified into two categories. The first category is Fourier Series-based analysis, which computes the Fourier decomposition of the intensity profile of the cell junctions (from 0° to 360° as a periodic signal) and determines polarity magnitude and angle using Fourier coefficients (Aigouy et al., 2010; Bardet et al., 2013; Merkel et al., 2014; Tetley et al., 2016; Banerjee et al., 2017; Aw et al., 2016). Although, this method has been widely used for 2D planar polarization quantification, however, in its current implementation the Fourier Series method shows significant sensitivity to cell geometry (see Results).

The second category calculates the ratio of fluorescence intensity of vertical cell edges to the horizontal cell edges as a readout of the polarity magnitude (Bulgakova et al., 2013; Farrell et al., 2017). As compared to the laborious manual classification of cell edges (Brittle et al., 2012; Ambegaonkar et al., 2012), this method automatically classifies the orientation of cell edges into both horizontal and vertical edges, with a prior assumption that the asymmetry of proteins is on either of these edges. Although this method has proved to be useful in some cases, it is poorly suited for cells that are irregular in geometry or for junctional proteins that are not polarizing along the vertical or horizontal axes of the cell. A variant of this method fits a square wave onto the junctional intensity profile and computes the ratio of opposite quadrants to determine the polarity magnitude and angle on a cell-by-cell basis (Strutt et al., 2016). Hence, this approach can be applicable to polarized junctional proteins on any cell axes, not limited to vertical or horizontal axes of the cell itself. However, due to the nature of this approach in fitting a square wave onto the junctional intensity profile, this approach is useful for regular shaped-cells but not irregular cells.

In this paper, we develop an unbiased and automated method to quantify the asymmetric distribution of proteins on cell boundaries based on Principal Component Analysis (PCA). This method computes the angle that produces the largest variance of weighted intensities from the centroid of the cell, which corresponds to the first principal component axis, and outputs the magnitude and angle of polarity, independently of cell geometries. To evaluate the efficacy and robustness of this method, we compare and validate this method against two other published methods (Fourier Series and Ratio methods) on both simulated models and experimental data. Additionally, we provide a user-friendly QuantifyPolarity Graphical User Interface (GUI) as a general tool for the study of epithelial tissue dynamics, such as quantification of planar polarization and several characteristics of cell morphology and topology (relationship with neighbors), allowing correlations between different cell properties to be explored. Together these provide powerful tools to understand how molecular and cellular mechanisms shape tissues.

## Results

### Validation of planar polarity quantification methods

There are two readouts of cell polarity: the strength of polarization (denoted as polarity magnitude) and the axis of polarization (denoted as polarity angle). If proteins are homogeneously distributed on the cell junctions, then the cell is considered to be unpolarized and polarity magnitude is zero for both the Fourier Series and PCA method and one for the Ratio method. However, if the proteins are asymmetrically segregated to opposite cell junctions, then the cell is considered to be polarized and has higher polarity magnitude. Polarity angle is defined as the axis that gives the maximum asymmetry.

As cells come in different shapes and sizes, which change throughout development, it is crucial to have an unbiased polarity quantification method that is unaffected by different cell sizes and shapes. For instance, larger cells should not be identified as more polarized when compared to smaller cells and more elongated cells should not be less polarized as compared to less elongated cells. Similarly, polarity angle should be oriented at the direction of maximum asymmetry, without being affected by different cell geometries. Here, we explore the sensitivity and robustness of different polarity methods in detecting varying strengths of polarization when challenged with varying cell sizes, shapes and eccentricities. Specifically, we validate the PCA method and compare it with two other established tools, namely the Ratio and Fourier Series methods, on simulated cells and images of protein distributions in different planar polarized tissues.

### Validation on simulated cells

To evaluate the robustness and accuracy of particular methods in the face of varying cell geometry, we simulated cells with varying size, shape regularity, eccentricity, and the amount of proteins on cell junctions.

Prior to *Drosophila* pupal wing hair formation, the Fz planar polarity protein becomes highly concentrated to distal cell junctions (Strutt, 2001) (Figure 1D). Due to the limited resolution of confocal microscopy, the localization signal of Fz on one side of a cell junction is inseparable from that on the other side, so Fz appears to be localized to both distal and proximal cell ends. Thus, from the junctional protein distribution profile, one expects two peaks of Fz protein intensity at *θ* and *θ* + *π*, which corresponds to the polarity angle (Figure 1D). For simplicity, we simulated a regular hexagonal cell with junctional proteins distributed on both horizontal and vertical junctions as illustrated in Figure 1D’. Each unit of junctional protein has an intensity value. Proteins on vertical junctions exhibit higher intensity (referred to as peak protein) while proteins on horizontal junctions exhibit lower intensity (referred to as base protein) (Figure 1D’). When we gradually increased the absolute amount of peak and base protein intensities (while maintaining the relative amounts) in the simulated cells, neither the polarity magnitude nor the angle is affected for all methods (Figure S1A-A’). Thus, the peak and base protein intensities were set to 255 and 40 arbitrary units (a.u.) respectively for all simulations. We then quantified polarity magnitude and angle obtained from different methods as a measure of how well simulated cells of different geometries were polarized. Polarity magnitudes obtained from different methods are normalized against their maximum magnitudes to allow direct comparison.

In *Drosophila* wild-type pupal wing cells, we determined that the average apical cell size varies between 2000 pixel^2^ to 2800 pixel^2^ when using our typical imaging settings (approximately 12 to 18 μm^2^) between 24 to 36 hours After Puparium Formation (hAPF) (Figure S1B). We attempted to further define the possible range of apical cell areas based on mutant data. Removing *dumpy* activity results in a shorter wing blade and reduced cell area due to a lack of extracellular matrix connections to the wing margin that counteract wing-hinge contraction (Etournay et al., 2015; Ray et al., 2015). The average apical cell area in *dumpy^ov1^* cells is significantly smaller than wild-type cells, being approximately 1700 pixel^2^ (approximately 10 μm^2^) at 30 hAPF (Figure S1E and E’). On the other hand, *ultrahair (ult)* and *cdc2* mutations in the *Drosophila* pupal wing produce substantially larger apical cell areas, about 4 to 16 times the size of normal cells (Adler et al., 2000). Hence, we simulated cells with apical areas ranging from approximately 1500 to 46000 pixel^2^ (approximately 10 to 300 μm^2^). Despite having different cell areas, these simulated cells had equivalent junctional protein distribution profiles (Figure 1E). Hence, these cells should have similar polarity magnitude and angle, and this is reflected in the outputs of all three polarity methods (Figure 1E and E’).

During *Drosophila* pupal wing development, cell shape changes from irregular to highly regular in geometry, resulting in an increasing fraction of regular hexagonal cells prior to wing hair formation (Classen et al., 2005) (Figure S1C). Loss of Rap1 results in highly aberrant cell shape as compared to wild-type cells (Knox and Brown, 2002). The average cell shape regularity in cells lacking Rap1 is approximately 0.6 as compared to a value of about 0.8 for wild-type cells at 30 hAPF (the value of cell regularity ranges between 0 and 1 with 0 being irregular and 1 being perfectly regular) (Figure S1E and E’’). Hence, we simulated cells with varying shape regularity value, ranging from 0.5 to 0.85, and fixed the amount of proteins (with peak proteins spanning approximately ±30° from the horizontal axis of an image) on the vertical cell junctions. We found that the polarity magnitudes and angles obtained from all methods are not affected by variation in cell regularities (Figure 1F-F’). These results suggest that all the polarity methods are suitable for quantifying planar polarity in cells with varying cell sizes and shapes.

Furthermore, (Aigouy et al., 2010) showed that *Drosophila* pupal wing cell elongation or eccentricity gradually decreases prior to wing hair formation. Indeed, from 24 to 36 hAPF, the average cell eccentricity of wild-type pupal wings decreases from 0.6 to 0.2 (the value of cell eccentricity ranges from 0 to 1 with 0 being circular and 1 being highly elongated) (Figure S1D). Hence, it is important to have a robust method for quantifying polarity that is not affected by different cell eccentricities. To examine if these polarity methods satisfy this criterion, we simulated cells with varying degrees of eccentricity (from 0 to 0.9) while maintaining peak and base protein intensities on both vertical and horizontal junctions respectively (Figure 1G-G’). In our simulations, the polarity magnitude computed using the PCA method is independent of varying degrees of cell eccentricity (Figure 1G). However, polarity magnitude obtained from both the Ratio and Fourier Series methods is sensitive to varying cell eccentricities. Our results showed that the Fourier Series method gave maximum polarity magnitude for cells with eccentricity of 0.7. Meanwhile, polarity magnitude computed using the Ratio method is significantly reduced for cells with eccentricity above and below 0.5. In terms of polarity angle measurement, all the methods give a constant polarity angle readout at 0° (Figure 1G’).

During *Drosophila* pupal wing development, Fz becomes increasingly polarized onto vertical junctions prior to wing hair formation before gradually depolarizing after the emergence of wing hairs (Usui et al., 1999; Classen et al., 2005; Aigouy et al., 2010; Merkel et al., 2014). Hence, we evaluated the performance of different methods in detecting different degrees of polarization strength. We generated cells with varying protein distribution profiles while maintaining their shape and sizes. First, we simulated a regular hexagon with a perimeter length of 440 units initially all set to a base protein intensity (40 a.u.). Next, we gradually increased the region of peak protein intensity (255 a.u.) on both vertical junctions starting at the poles of the cell and moving onto the horizontal junctions as illustrated in Figure 1H’’.

As predicted, an initial simulated cell with an homogenous distribution of base proteins exhibited no polarization (Figure 1H and H’’). Gradual increments of junctional peak proteins on the poles of both the vertical junctions results in increasing polarity magnitude, however, the maximum polarity magnitude obtained from all the three methods varied. Maximum polarity is achieved when the amount of the cell perimeter comprising junctional peak protein is approximately 200 units for Ratio, 160 for Fourier Series and 240 for PCA as indicated by the arrows in Figure 1H and H’’. Polarity magnitudes then decrease steadily and subsequently drop to zero when junctional peak protein is homogenously distributed on the entire cell perimeter (Figure 1H and H’’). In terms of polarity angle, all methods remain consistently oriented at 0° despite varying protein distribution (Figure 1H’). From this simulation, we found that there is a distinct polarization strength profile for each of the methods. This allows users to choose a method which gives them an appropriate profile accordingly to their requirements. For example, prior to *Drosophila* pupal wing hair formation, Fz becomes highly polarized to the distal cell junctions (Strutt, 2001). This is represented by simulated cell with 320 units of peak proteins equally segregated to opposite vertical junctions (Figure 1H’’). Thus, the PCA method, which gives maximum polarity magnitude at approximately 240 units of peak protein, may be best suited to quantify polarization in pupal wings.

Taking together these simulation results, the PCA method is the only method that successfully quantifies polarity in an unbiased manner, independently of different cell size, shape and eccentricity. However, each polarity method has its own unique polarity strength profile, which could be advantageous for the analysis of different types of polarized cells.

### Validation of different polarity quantification methods on biological datasets

Although the PCA method that we developed performs rather effectively in quantifying polarity on simulated cells, this might not necessarily reflect its performance on biological datasets. We therefore compared results obtained from the PCA method with the Fourier Series and Ratio methods, on images of different planar polarized epithelial tissues, specifically the *Drosophila* pupal wing, third instar wing discs and the embryonic epidermis, which each exhibit distinct cell geometries (Figure 2A-B).

**Figure 2.**
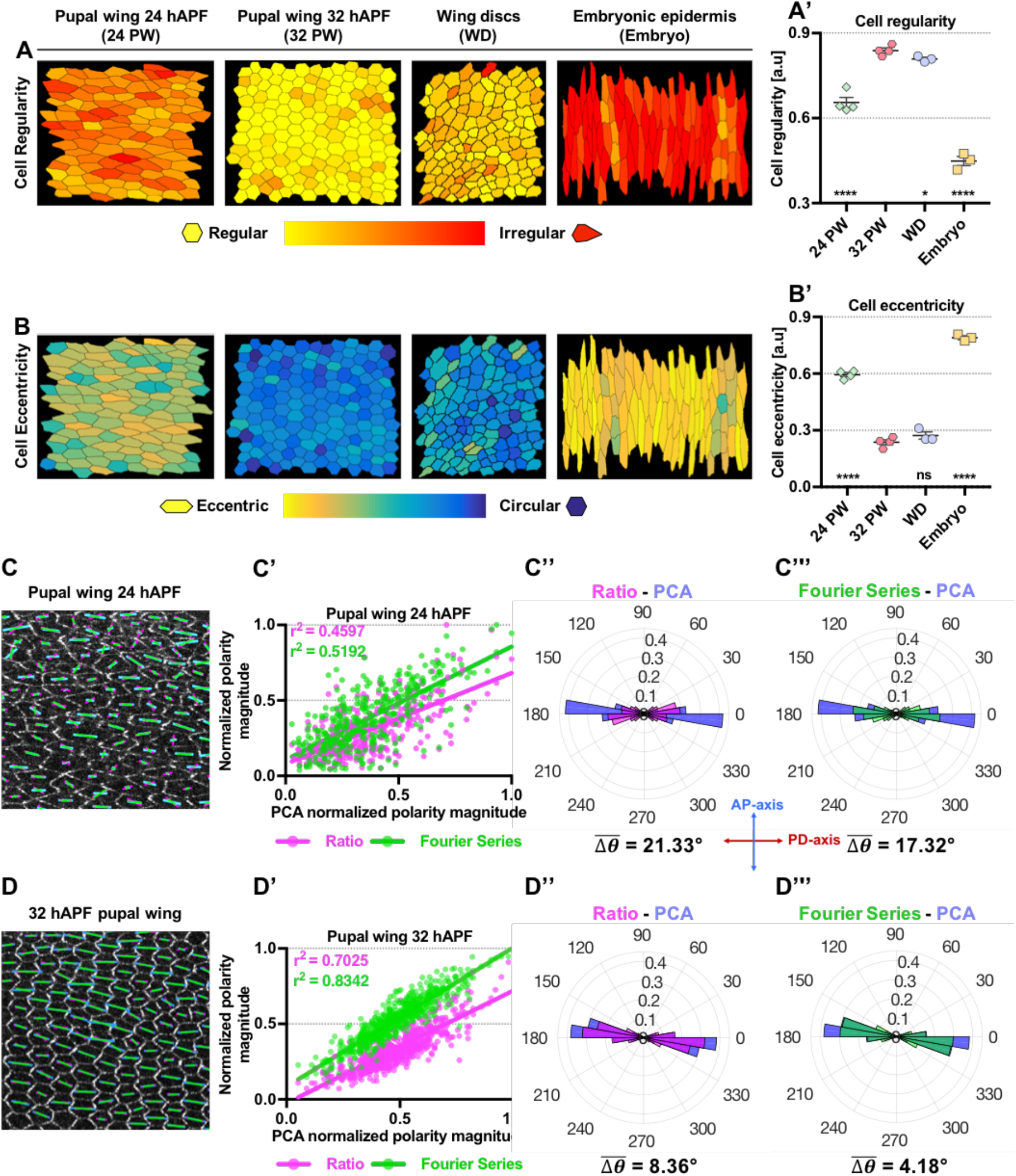
Comparison of different methods for planar polarity quantification on *Drosophila* pupal wings at different developmental time points. (A-B) Processed images of various planar polarized epithelial tissues: 24 and 32 hAPF *Drosophila* pupal wing, third instar wing discs and the embryonic epidermis, which each exhibit different cell regularity and eccentricity as compared to regularly packed hexagonal cells. (A-A’) Cells are color-coded according to the regularity of the shape, with yellow being perfectly regular and red represent highly irregular. (A’) Quantified average cell shape regularity of 24 and 32 hAPF pupal wings, wing discs and embryonic epidermis. (B-B’) Cells are color-coded according to the eccentricity of the shape, with yellow represent highly eccentric and blue being circular or non-eccentric. (B’) Quantified average cell eccentricity of 24 and 32 hAPF pupal wings, wing discs and embryonic epidermis. (C and F) Quantified cell-scale polarity pattern of otherwise wild-type wings expressing Frizzled-EGFP at (C) 24 hAPF and (F) 32 hAPF using three different methods. The blue, green, magenta bars represent the magnitude (length of bar) and angle (orientation of bar) of planar polarization pattern for a given cell obtained from PCA, Fourier Series and Ratio methods respectively. (D) Plot of normalized polarity magnitudes at 24 hAPF obtained from Ratio (magenta dots) and Fourier Series (green dots) versus PCA methods with best fit lines shown as magenta and green lines respectively. Coefficients of determination r^2^ are indicated. (E-E’) Circular weighted histogram plots displaying the orientation of Fz-EGFP polarity obtained from (E) Ratio and PCA and (E’) Fourier Series and PCA methods at 24 hAPF with mean angle differences indicated respectively. (G) Plot of normalized polarity magnitudes at 32 hAPF obtained from Ratio (magenta dots) and Fourier Series (green dots) versus PCA methods with best fit lines shown as magenta and green lines respectively. Coefficients of determination r^2^ are indicated. (H-H’) Circular weighted histogram plots display the orientation of Fz-EGFP polarity obtained from (H) Ratio and PCA and (H’) Fourier Series and PCA methods at 32 hAPF with mean angle differences indicated respectively. In (A’-B’) the number of wings per genotype is 3 to 4. Error bars indicate mean±SEM. One-way ANOVA unpaired test, comparing each genotype to 32 hAPF pupal wing. Significance levels: p-value ≤ 0.0001^****^, p-value ≤ 0.01^*^ and ns is not significantly different. Polarity magnitudes in (D and G) obtained using each different method are normalized to allow comparison. Polarity angles (in degree) in (E-E’ and H-H’) range between 0° to 360°, with 0° corresponding to the x-axis of the image.

### *Drosophila* pupal wing analysis

The simulation results allow us to better understand the behavior of each polarity method on either varying cell morphology or protein distribution. However, none of the simulated cases above represent the real biological scenario, in which both cell morphology and protein distribution change in concert. This is particularly striking during late *Drosophila* pupal wing morphogenesis, where cell polarity intensifies as cell shape changes from irregular to regular geometry (Classen et al., 2005; Aigouy et al., 2010). Apart from changes in cell shape regularity, *Drosophila* pupal wing epithelial cells also undergo significant changes in apical area, eccentricity and orientation over these developmental time points (Figure S1 and S4) (Aigouy et al., 2010). Hence, measuring polarity during late *Drosophila* pupal wing morphogenesis provides a dynamic system for comparing the performance of different methods on biological data. We therefore analyzed the correlation between the polarization magnitude obtained from the different methods of EGFP-tagged Fz (a core planar polarity protein) in otherwise wild-type wings at two developmental stages (Figure 2C-F). We found that at 24 hAPF when cells are more eccentric there is only a moderate correlation in polarity magnitude obtained from the Ratio (coefficient of determination, r^2^ = 0.4597) and Fourier Series (r^2^ = 0.5192) with the PCA method (Figure 2D). This moderate degree of correlation is likely to be due to the fact that polarity magnitudes computed from the Fourier Series and Ratio methods are more sensitive to cell eccentricity, as evident from the simulation results (Figure 1G). However, the correlation between the Ratio and Fourier Series to PCA method improved (with r^2^ = 0.7025 and 0.8342 respectively) as cells become less eccentric and more regular by 32 hAPF (Figure 2G).

We computed the mean angle difference, 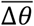, to compare polarity angles obtained from the PCA against the Ratio and Fourier Series methods (See “Circular weighted histogram” in Materials and Methods). As compared to 32 hAPF, polarity angles obtained from both the Ratio and Fourier Series methods at 24 hAPF are less in agreement with the PCA method (with Δ*θ* of 21.33° and 17.32° respectively) (Figure 2E-E’). However, by 32 hAPF, polarity angles computed from both methods agree better with the PCA method (with mean angle differences of 8.36° and 4.18° respectively) (Figure 2H-H’). Note that earlier in development at 24 hAPF, polarity angles are more dispersed as compared to 32 hAPF (with angle variance of 0.137±0.013 and 0.059±0.0055 at 24 and 32 hAPF) where the polarity pattern becomes more coherent and well-aligned along the proximal-distal axis (PD-axis) (Figure 2E-E’ and 2H-H’).

As a comparison to the polarized distribution of Fz-EGFP, we also quantified the polarization of E-Cadherin::GFP at 32 hAPF, which shows only weak asymmetry in the pupal wing (Warrington et al., 2013) (Figure S2A). Quantification of E-Cadherin::GFP distribution with all the polarity methods gives low but non-zero polarity magnitude (Figure S2A-B). Furthermore, E-Cadherin::GFP distribution also exhibits dispersed angles (Figure S2C-C’). To capture the local coordination of polarity within a group of cells, we quantified and compared the coarse-grain polarity of E-Cadherin::GFP and Fz-EGFP at 32 hAPF. Indeed, vector average polarization of E-Cadherin::GFP is much smaller than Fz-EGFP (Figure S2D’), with E-Cadherin::GFP showing at best very weak local polarity coordination (Figure S2D).

Note that even for unpolarized (but non-homogenous e.g. punctate) distribution of proteins, each method nevertheless reports a readout of polarity magnitude and angle. We therefore suggest users should use polarity magnitude as a weight for its angle in a circular histogram as in e.g. Figure 2E-E’ (see “Circular weighted histogram” in Materials and Methods), rather than simply plotting unweighted polarity angles (Aw et al., 2016). Hence, quantifying a poorly polarized protein such as E-Cadherin provides a baseline polarity readout, which can be an important control for comparison with well-polarized proteins.

### *Drosophila* wing discs

Next, we quantified the asymmetric localization of the Dachsous planar polarity protein in third-instar larval wing imaginal discs using all three polarity methods (Figure 3A). In terms of polarity magnitude, both the Ratio and Fourier Series methods seem to correlate well with the PCA method (r^2^ = 0.7195 and 0.7687) (Figure 3B). Moreover, polarity angles obtained from both the Ratio and Fourier Series methods are not significantly different from the PCA method (with 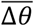 of 14.59° and 10.63° respectively) (Figure 3C-C’). In fact, there is only a subtle difference between the geometry of third-instar central wing pouch cells from that of 32 hAPF pupal wing cells (Figure 2A’-B’), and so similarly there is good correlation between all three polarity methods.

**Figure 3.**
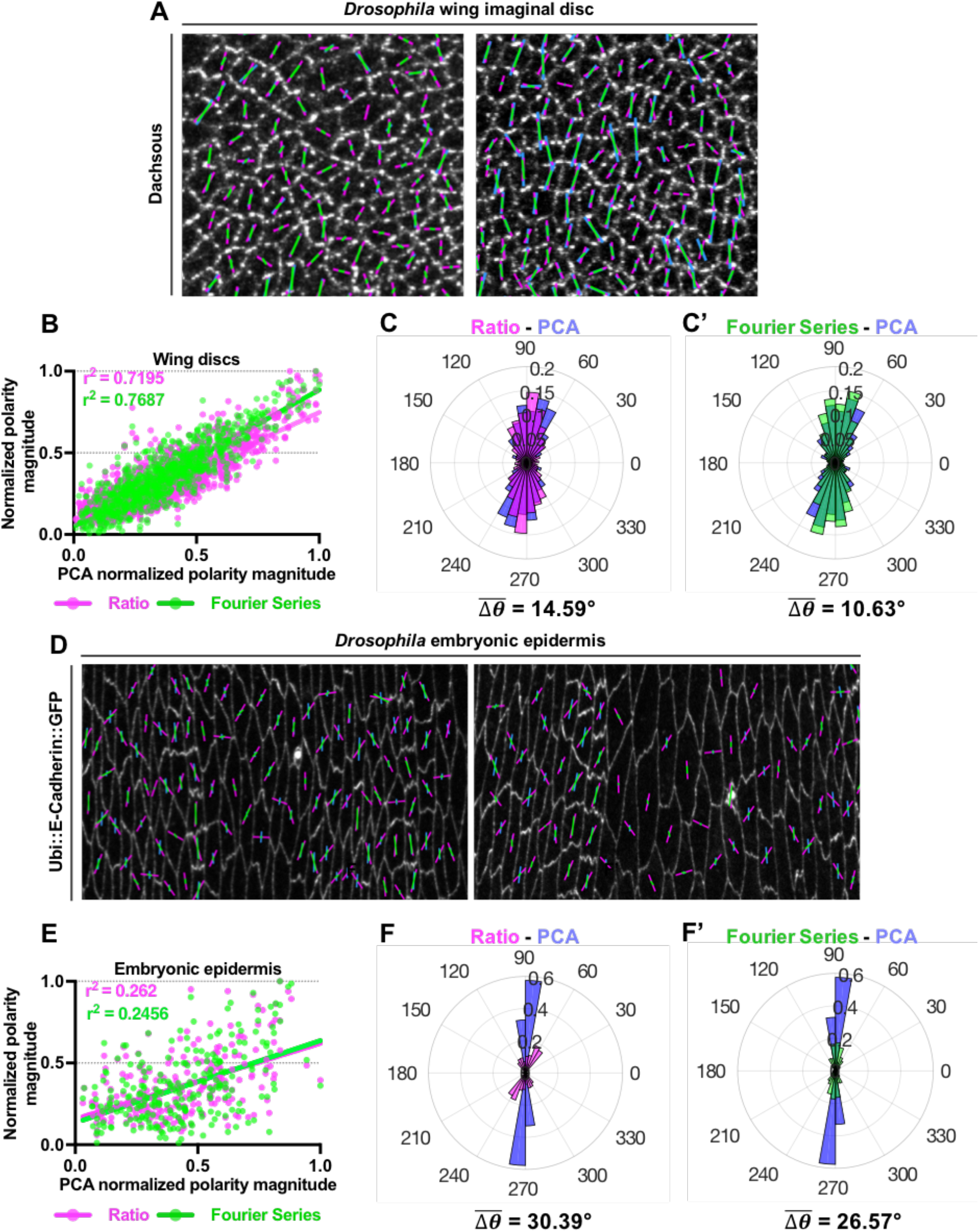
Validation of different methods for quantification of planar polarity on *Drosophila* wing imaginal discs and embryonic epidermal cells. (A) Quantified cell-scale polarity pattern of two examples of wing discs immunolabeled for Dachsous in third-instar larval imaginal discs. The magenta, green and blue bars represent the magnitude and orientation of planar polarity for a given cell obtained from Ratio, Fourier Series and PCA methods respectively. (B) Plot of normalized polarity magnitudes obtained from Ratio (magenta dots) and Fourier Series (green dots) versus PCA methods with best fit lines shown as magenta and green lines respectively. Coefficients of determination r^2^ are indicated. (C-C’) Circular weighted histogram plots display the orientation of Dachsous polarity obtained from (C) Ratio and PCA and (C’) Fourier Series and PCA methods with its mean angle differences respectively (*n* = 3 wing discs with 900 cells analyzed). (D) Two examples of quantified cell-scale polarity pattern of Ubi::E-Cadherin-GFP expressing epidermal embryonic cells at stage 15. Dorsal is to the top of these images. (E) Plot of normalized polarity magnitudes of embryonic epidermal obtained from Ratio (magenta dots) and Fourier Series (green dots) versus PCA methods with best fit lines shown as magenta and green lines respectively. Coefficients of determination r^2^ are indicated. (F-F’) Circular weighted histogram plots display the orientation of Ubi::E-Cadherin::GFP polarity obtained from (C) Ratio and PCA and (C’) Fourier Series and PCA with its mean angle differences respectively (*n* = 3 embryos with 250 cells analyzed). In panels B and E, polarity magnitudes (a.u) obtained using each different method are normalized accordingly. All polarity angles (in degree) range between 0° to 360°, with 0° corresponding to the x-axis of the image.

### *Drosophila* embryonic epidermis

Lastly, we quantified Ubi::E-Cadherin-GFP asymmetry in images of lateral epidermal cells in *Drosophila* embryos at stage 15, where the embryonic epidermal cells exhibit an elongated rectangular shape (Figure 3D). The embryonic epidermal cells are much more irregular and eccentric in geometry as compared to both 24 and 32 hAPF pupal wing and wing disc cells (Figure 2A’-B’). Based on published results from (Bulgakova et al., 2013), E-Cadherin is asymmetrically localized to the shorter cell boundaries (dorsal-ventral), therefore should exhibit an approximately ±90° angle of polarization (along the y-axis in the image). Notably, polarity angles of embryonic epidermal cells computed using the PCA method are well-aligned along ±90° (with angle variance of 0.038±0.013), in agreement with the published results (Bulgakova et al., 2013) (Figure 3F). On the contrary, polarity angles obtained from both the Ratio and Fourier Series methods are more dispersed from −90° to +90° (with angle variances of 0.52±0.04 and 0.38±0.03 respectively), which disagrees with the previous analysis (Bulgakova et al., 2013) (Figure 3F-F’). Overall, the polarity angles obtained from both the Ratio and Fourier Series methods are less aligned with the PCA method (with 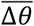 of 30.39° and 26.57° respectively) (Figure 3F-F’). These results showed that the accuracy of both the Fourier Series and Ratio methods in detecting polarity angle are affected by elongated cells. In terms of polarity magnitude, both the Ratio and Fourier Series methods are also poorly correlated with the PCA method (r^2^ = 0.262 and 0.2456 for Ratio and Fourier Series respectively) (Figure 3E). As the epidermal embryonic tissues present with a mixture of both moderately-elongated and highly-elongated cells, polarity magnitudes computed from the Fourier Series and Ratio methods will be affected by variation in cell eccentricities, as evident from the simulation results (Figure 1G).

### QuantifyPolarity Graphical User Interface (GUI): an automated tool for quantification of planar polarization and cell shape

The QuantifyPolarity Graphical User Interface (Figure 4) was developed to provide a tool for fast and reliable analysis of 2D planar polarity in multicellular tissues by incorporating all the three quantification methods described here – Ratio, Fourier Series and PCA. These methods are applicable to any 2D asymmetrical distribution of proteins on cell junctions, for example, asymmetrical localization of auxin-efflux carrier PIN1 in leaf primordia or polarization of Myosin II during germband extension (Benková et al., 2003; Zallen and Wieschaus, 2004; Kasza et al., 2014; Tetley et al., 2016). Generally, all three methods are able to accurately quantify planar polarization on cells with regular geometry, however, for highly irregular or elongated cells, PCA method may be a better choice to provide unbiased results.

**Figure 4.**
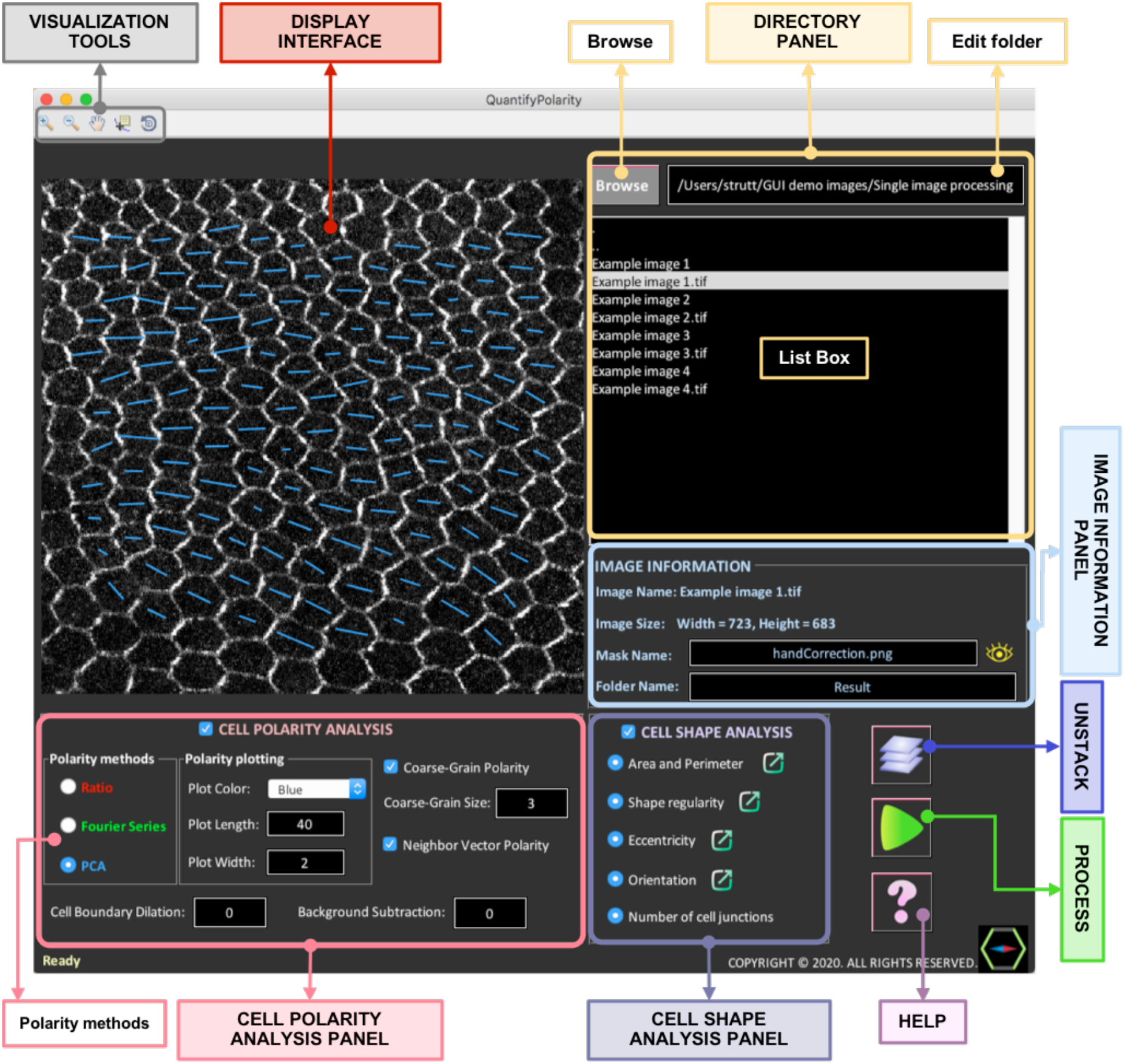
An overview of all the components available in the QuantifyPolarity Graphical User Interface. The QuantifyPolarity GUI consists of six interactive panels: Visualization tools, Display interface, Directory panel, Image information panel, Cell polarity analysis panel and Cell shape analysis panel.

The cell-by-cell polarity readout obtained from the Ratio, Fourier Series and PCA methods reveals the polarization strength (magnitude) and alignment (angle) of each individual cell (Figure S3B’(i)). Averaging this value (“Average Polarity Magnitude”) gives a measure of polarization strength of all cells within the image, without taking into consideration the coordination of polarity between cells. To provide a combined measure that takes into account both polarity magnitude and polarity coordination over a group of cells, we calculate the vector average of individual cell-by-cell polarity magnitudes and angles (see Materials and Methods). When applied over the whole image, we refer to this measure as the “Vector Average Polarity Magnitude”, when applied to a cell and its immediate neighbors we call it the “Average Neighbor Vector Polarity Magnitude” and when applied over a local group of cells we call it the “Coarse-Grain Vector Polarity” (Figure S3B’(ii-iii)). The “Angle Variance” measures the variance in polarity alignment of all cells within the image. See Figure 7 for an explanation of the differences between each polarity measurement on various examples of polarized tissues.

In addition to polarity quantification, QuantifyPolarity also includes quantitative analysis of several cell morphological properties (e.g. size, shape regularity, eccentricity and orientation) and topology (number of neighbors), which are useful for the study of morphogenesis (Figure S3B’’). QuantifyPolarity also generates (customizable) color-coded images corresponding to the quantitative measurements, allowing users to directly visualize and inspect the results of the quantification. For example, each cell is color-coded with a gradient color-map according to their apical area, shape regularity, eccentricity and number of cell junctions (Figure S4) allowing visualization of the temporal and spatial evolution of cell geometries. Additionally, all results generated by QuantifyPolarity are automatically organized into Comma-Separated Values (CSV) files for easy accessibility and further analysis.

Since cell dynamics could evolve differently in different regions of the tissue, or mutant cells could behave differently as compared to neighboring control cells, we added a feature which allows the user to perform multiple analysis (for e.g. different polarity methods, different ROIs and so on) on the same image. This feature allows the user to analyze and compare the cell dynamics within different ROIs. Equipped with the functionality of batch processing, users can automate and accelerate the analysis of multiple images within the same folder, which is often a time-consuming and tedious process. Finally, QuantifyPolarity can operate on wide range of platforms such as Mac and Windows without requiring additional software.

### Analysis of temporal evolution of planar polarity and cell morphology in QuantifyPolarity

As a demonstration of the functionalities of QuantifyPolarity, we used it to investigate the temporal evolution of cell polarity and cell morphological properties during *Drosophila* pupal wing development. We quantified polarization magnitude of Fz-EGFP in the proximal-posterior region of otherwise wild-type wings (Figure 5A) from 24 to 36 hAPF using the three polarity methods (Figure 5B-B’).

**Figure 5.**
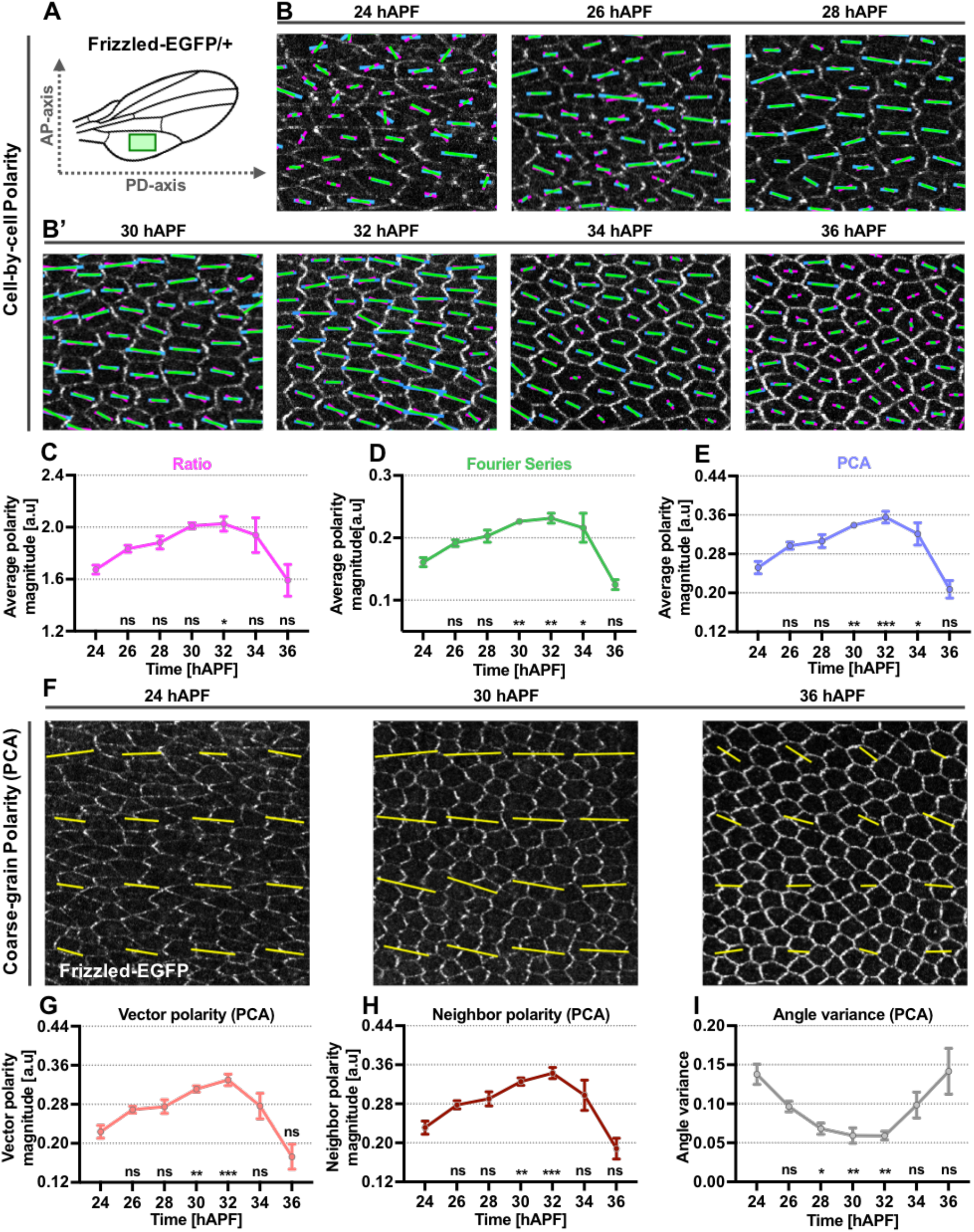
Application of different methods for planar polarity quantification on *Drosophila* pupal wings at different developmental time points. (A) Illustration of analyzed proximal-posterior region below vein 5 of the wild-type pupal wing blade (represented by a green box). (B-B’) Quantified cell-scale polarity pattern of otherwise wild-type wings expressing Fz-EGFP from 24 hAPF to 36 hAPF. The magenta, green and blue bars represent the magnitude (length of bar) and angle (orientation of bar) of planar polarization for a given cell obtained from the Ratio, Fourier Series and PCA methods respectively. (C-E) Plots of average polarity magnitudes (in a.u.) obtained from the Ratio (C), Fourier Series (D) and PCA methods (E) respectively for Fz-EGFP wings at the indicated time points. (F) Quantified coarse-grain polarity pattern of otherwise wild-type wings expressing Fz-EGFP at 24, 30 and 36 hAPF. The yellow bars represent the magnitude (length of bar) and angle (orientation of bar) of planar polarization for a group of cells obtained from the PCA method. (G-I) Plots of (G) vector average polarity magnitude, (H) neighbor average polarity magnitude and (I) angle variance obtained from the PCA method for Fz-EGFP wings at the indicated time points. The number of wings for each developmental time point is between 4 to 5. Error bars indicate mean±SEM. One-way ANOVA unpaired test, comparing each time point to 24 hAPF pupal wing. Significance levels: p-value ≤ 0.0001^****^, p-value ≤ 0.0005^***^, p-value ≤ 0.002^**^, p-value ≤ 0.01^*^. ns is not significantly different.

All three methods displayed a similar trend in which average polarity magnitude gradually increases from 24 to 32 hAPF and then decreases from 32 to 36 hAPF (Figure 5C-E). Note depolarization of core polarity protein occurs following the formation of actin-rich trichomes at 32 hAPF (Usui et al., 1999; Merkel et al., 2014). By comparing each developmental time point to 24 hAPF, we found that the PCA method gave higher statistical significance in detecting overall changes in polarity distribution as compared to the other methods (Figure 5C-E). In support of this, we also computed the one-way ANOVA F-ratio which is the ratio of variability between average polarity for each time point and variability within the time point for a given polarity method. The F-ratio obtained from the PCA method is higher as compared to both the Ratio and Fourier Series methods, indicating that there is a higher statistical significance between average polarity magnitude for each time point (F-ratios are approximately 4.7 for Ratio, 12.7 for Fourier Series and 15.9 for PCA). This could be attributed to the differences in polarization strength profile for different methods: Based on our simulation results, both the Fourier Series and Ratio methods attained maximum polarization magnitude even when proteins are not fully segregated to the opposite vertical junctions (Figure 1H’). Moreover, both the Fourier Series and Ratio methods are affected by variation in cell eccentricities, reporting lower polarity magnitude for less eccentric cells and attained maximum polarity magnitude at cell eccentricity around 0.5 for Ratio and 0.7 for Fourier Series (Figure 1G). As cells become more regular in shape from 24 to 36 hAPF, cell eccentricity decreases from approximately 0.6 to 0.2 (Figure S1D). This results in lower polarity magnitude readout at later time points, thereby reduces the differences in polarity strength between earlier and later time points. This suggests that the PCA method is more sensitive and reliable in detecting changes in protein distribution accompanied with variation in cell geometry.

Next, we used the polarity readout from the PCA method to perform a broader polarity analysis during *Drosophila* pupal wing morphogenesis. Both the PCA vector average and neighbor average polarity magnitudes gradually increased from 24 hAPF to 32 hAPF and then decreased from 32 to 36 hAPF, displaying the same trend as the average polarity magnitude readout (Figure 5E, G and H). However, average polarity strength in 34 hAPF pupal wings was significantly greater than 24 hAPF pupal wings (Figure 5E), while this difference is no longer significant when considering the vector and neighbor average polarity (Figure 5G-H). This is due to these vectorial measures capturing both polarity strength and local polarity alignment, with the latter being low at both earlier and later timepoints. This is reflected in the polarity angle variance which decreases from 24 to 32 hAPF as polarity alignment increases, then decreases from 32 to 36 hAPF as polarity angles become more dispersed (Figure 5I).

The pioneering works from (Classen et al., 2005; Aigouy et al., 2010) have demonstrated that prior to emergence of polarized wing hairs, the magnitude of core planar polarization intensifies concomitantly with relaxation of cells into a regular hexagonally packed geometry. It was also reported that core planar polarity magnitude is temporally correlated with the magnitude of cell elongation during pupal wing morphogenesis (Aigouy et al., 2010). To understand the mechanism by which epithelial tissues develop specific packing geometries and coordinate their core planar polarity, we examined how cell size and shapes correlate with the strength of core protein polarization from 24 to 32 hAPF of otherwise wild-type wings. Consistent with previous findings, the temporal progression of Fz-EGFP polarity magnitude strongly correlated with changes in cell regularity and eccentricity over these developmental time points. Average cell shape regularity and polarity magnitude were positively correlated (with coefficient of determination, r^2^ = 0.9116), in that more regular cells exhibited higher polarity magnitude and vice versa (Figure 6A). Similarly, average cell eccentricity and polarity magnitude were negatively correlated (with coefficient of determination, r^2^ = 0.9246), with less elongated cells being more polarized and vice versa (Figure 6B). Interestingly, we found only a weak correlation between apical cell area and polarity magnitude during these developmental time points in wild-type pupal wings (with coefficient of determination, r^2^ = 0.238) (Figure 6C).

**Figure 6.**
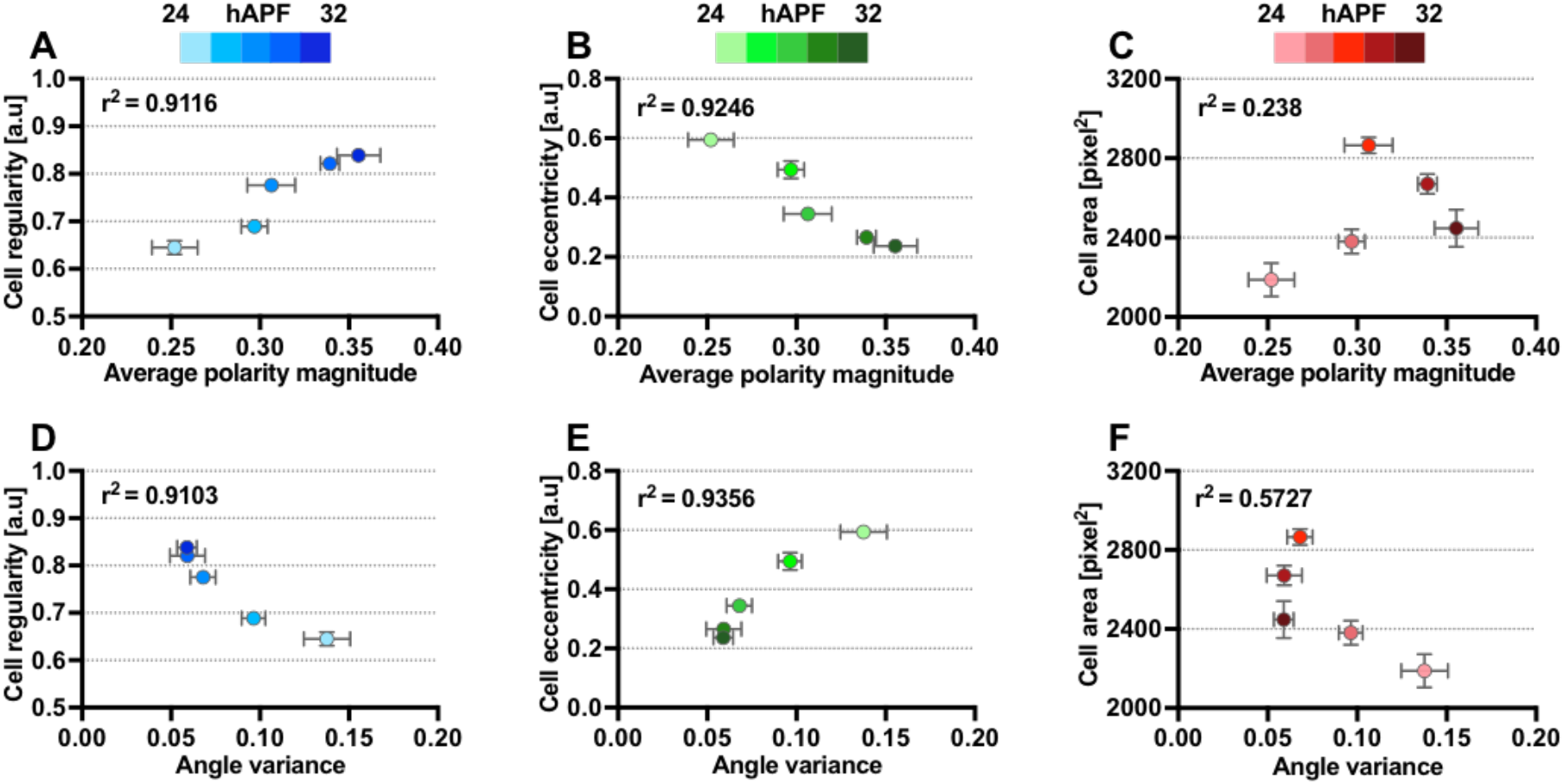
Temporal correlation between cell size, regularity and eccentricity with Fz-EGFP polarity of wild-type wings. (A) Positive correlation between average cell regularity and Fz-EGFP polarity magnitude. (B) Negative correlation between average cell eccentricity and Fz-EGFP polarity magnitude. (C) Lack of correlation between apical cell area and Fz-EGFP polarity magnitude. (D) Negative correlation between average cell regularity and Fz-EGFP polarity angle variance. (E) Positive correlation between average cell eccentricity and Fz-EGFP polarity angle variance. (F) Lack of correlation between apical cell area and Fz-EGFP polarity angle variance. Each dot represents the total average of averaged values from all wings for specific developmental time point. The number of wings for each developmental time point is between 4 to 5. Error bar indicates mean±SEM.

**Figure 7.**
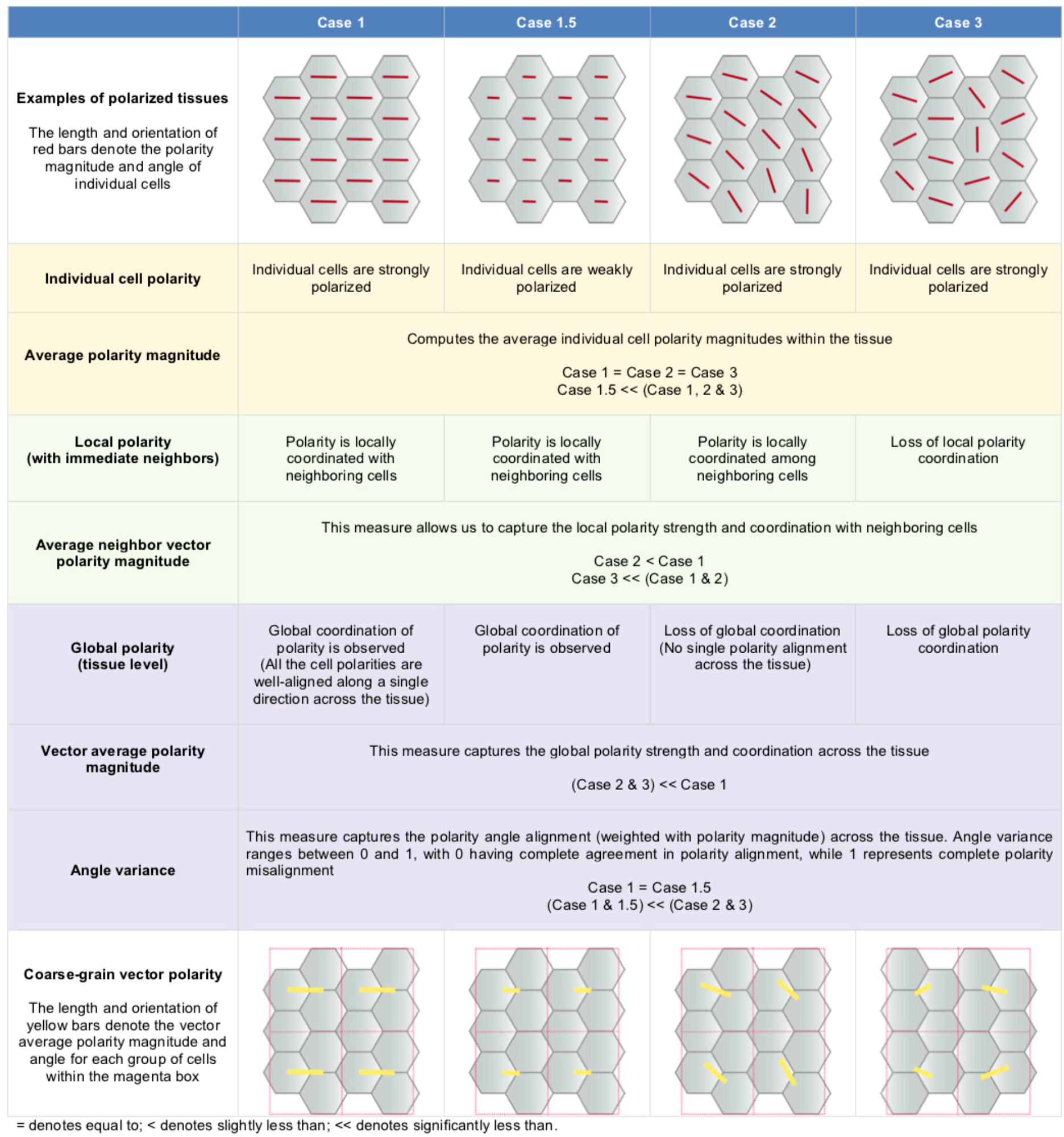
Five different polarity measurements to quantify polarization at different scales (cellular, local and global)

As wild-type wing tissue becomes more regularly packed and less eccentric, Fz reorients its polarity alignment to become increasingly coordinated along the PD-axis of the wing (Classen et al., 2005; Aigouy et al., 2010). Hence, we examined if cell size, regularity and cell eccentricity correlated with core polarity alignment. Indeed, we found that polarity angle variance was strongly correlated with cell regularity and eccentricity (with coefficient of determination, r^2^ = 0.9103 and 0.9356 respectively) but moderately correlated to apical cell size (with coefficient of determination, r^2^ = 0.5727) during late pupal development (Figure 6D-F). Thus, when using the PCA method as an accurate measure of planar polarization, we are able to conclude that Fz planar polarity does indeed increase as cell packing becomes more regular and cell eccentricity decreases. We furthermore find negligible evidence for cell size influencing planar polarity at this stage of pupal wing development.

## Discussion

Planar polarization is an essential process during morphogenesis for coordinating and organizing cells to establish specific tissues structures in a wide range of organisms. Hence, accurate and unbiased quantitative analysis of planar polarization is of paramount importance for deciphering molecular mechanisms underlying morphogenesis. Previous planar polarity quantification methods can be affected by variation in cell geometry. This study describes a novel method for quantifying asymmetrical localization of junctional proteins based on Principal Component Analysis (PCA). This method has been validated against existing polarity methods (the Fourier Series and Ratio methods) under various conditions using computer simulated cells. The simulation results revealed that the polarity readout from both the Fourier Series and Ratio methods are robust against variation in cell sizes and regularities but not cell eccentricities. The PCA method, on the other hand, consistently produces a polarity readout that is unaffected by variation of cell sizes and shapes. Furthermore, each polarity method exhibits a unique polarity strength profile to accommodate diverse types of polarized cells. Having validated these methods on simulated data, we tested the performance of these methods on various planar polarized epithelial tissues with distinctive cell geometries. Existing polarity methods correlate well with the PCA method on regular and less eccentric cell shapes. However, consistent with the simulation results, polarity readouts obtained from both the Fourier Series and Ratio methods are poorly correlated with the PCA method (and the published results) on highly elongated epidermal embryonic cells. Both simulation and experimental results demonstrate that the PCA method can be used reliably to quantify planar polarization independently of cell geometries.

To allow for automated and high-throughput analysis of cell polarity and shape, we further developed a standalone and user-friendly graphical user interface, QuantifyPolarity. This general tool enables experimentalists with no prior computational expertise to perform comprehensive analyses of cellular and molecular mechanisms driving tissue morphogenesis. To demonstrates the application of QuantifyPolarity, we analyzed the temporal dynamics of cell behavior in the developing pupal wing. Here we found that the temporal progression of core planar polarization magnitude is strongly correlated with cell regularity and eccentricity, consistent with a previous report (Aigouy et al., 2010). While it is clear that correlation does not necessarily imply causation, it will be interesting to investigate the causality effect of cell shape on core planar polarization. Although it is known that apical cell area plays a role in affecting core planar polarity system, where Fz fails to restrict prehair initiation to the distal cell junctions in substantially larger cells (Adler et al., 2000), there is a lack of temporal correlation between core planar polarization and apical cell size of otherwise wild-type wings. This is likely due to the fact that these apical cell sizes in wild-type wings fall within the “normal” range. Hence, it will be interesting to examine how considerably smaller or larger cell size affect the ability of core proteins to polarize.

Similarly, there is a strong temporal correlation between global polarity alignment and cell regularity and eccentricity but weaker correlation with apical cell area in wild-type wings. It has been proposed that irregular epithelial packing poses a challenge for feedback propagation of polarization signal across the epithelium (Ma et al., 2008) and defective hexagonal packing leads to loss of global polarity coordination in the wing (Bardet et al., 2013). However, it has been reported that stretch-induced directional cell junctional rearrangement plays a role in coordinating global polarity alignment (Aigouy et al., 2010; Aw et al., 2016). Thus, polarity alignment may not be simply a consequence of cell geometry. An understanding of how different cell geometry quantitatively accounts for the underlying mechanism of core planar polarization can serve as a route to understanding molecular mechanisms of tissue planar polarization.

## Materials and Methods

### Dissection and mounting of pupal wings for *in vivo* live imaging

All the fly strains used in this study are described in Supplementary Table 1 and were raised at 25°C. Pupae were dissected and mounted for *in vivo* live imaging as described in (Classen et al., 2008) as live imaging is less susceptible to potential artefacts (e.g. noise from non-specific labelling or changes in tissue shape due to dissection and fixation). Briefly, pupae were placed on a piece of double-sided tape dorsal side up. Using a pair of fine scissors and forceps, the puparium case was carefully removed from above the developing pupae to expose the wing without injuring the pupa. The exposed pupal wing was covered in a drop of Halocarbon 700 oil and was then taped onto a 2.5 cm glass-bottomed dish (Iwaki) with the wing facing the coverslip.

Subsequent imaging and processing steps are summarized in Figure S3 and described below.

### Preparation of wing discs and embryos for fixed imaging

Wing discs were immunolabeled for Dachsous protein distribution and imaged as described (Hale and Strutt, 2015). Embryos expressing Ubi::E-cadherin-GFP were fixed and imaged as described (Bulgakova et al., 2013).

### Image acquisition

Live image acquisition was performed using an inverted Nikon A1 confocal microscope with a Nikon 60x apochromatic objective lens oil (NA = 1.4) and GaAsP detectors. The pinhole was set to 1.2 Airy Unit (AU). A heated stage was set to 25°C. To maintain constant power for all imaging sessions, laser power was checked and if necessary adjusted before each imaging session. For imaging of green emissions, a 488 nm laser with a 525-550 band pass filter was used to detect EGFP. Images were taken at the posterior region of the pupal wing with 1024 × 1024 pixels per z-slice and 80 nm pixel size. For each wing, 12-bit z-stacks (with ~40 slices per stack, 0.5 μm/slice) were acquired. After time-lapse imaging, pupae were kept and survived to at least pharate stage and >95% to eclosion stage.

### Image processing

Raw microscopy images were first processed using external tools (e.g. PreMosa and PackingAnalyzer) to obtain skeletonized representation of the cell boundaries (also known as segmented images). These segmented images along with their original images are the pre-requisite inputs in QuantifyPolarity GUI for further image analysis.

### (i) Image surface extraction (PreMosa)

For image processing, microscopy images were exported into .TIFF format using Nikon software (NIS-Elements AR) for further processing. These z-stack images were automatically surface extracted and projected using PreMosa as described in (Blasse et al., 2017) to obtain a 2D projected image of the apical band of monolayer epithelial tissues (Figure S3A). To quantify proteins localizing to the apical junctions, Fz-EGFP z-stack images were used to generate the height map. In brief, this algorithm generates an initial height map that contains information of each z-slice with the brightest pixels. To yield a smooth and optimize height map, smoothing (with a median filter) and artefact correction processes were carried out. The final height map was used to project the manifold of interest onto a 2D image.

More commonly used image surface extraction method including maximum intensity projection (available in Fiji). Note that this step can be omitted for single z-slice image acquisition.

### (ii) Image segmentation (PackingAnalyzer)

To identify epithelial cell boundaries, the image was segmented using the semi-automated cell segmentation software, PackingAnalyzer (Aigouy et al., 2010). Cell segmentation software such as PackingAnalyzer is utilized to identify cell boundaries using a watershed algorithm (Aigouy et al., 2010). This procedure semi-automatically identifies and produces a binary skeletonized representation of the cell boundaries for further image analysis (Figure S3A). Additional manual correction was often required to obtain precise segmentation of cell boundaries. Thus, all segmented images were checked and corrected manually for segmentation errors such as under-segmentation and over-segmentation. Boundary cells and small cells are automatically removed based on the area thresholds set by users, which vary according to the image size and specifications. This is because boundary cells do not contain all the cell edges, therefore, it is not possible to quantify the morphological properties of boundary cells. These segmented images are then passed onto the QuantifyPolarity GUI for further image analysis.

Other useful image segmentation tools are SEGGA, EpiTools and TissueAnalyzer (newer version of PackingAnalyzer available as a Fiji plugin) (Farrell et al., 2017; Heller et al., 2016; Etournay et al., 2016).

### Identification of cells and neighbor relations

A series of steps were then employed to extract information out of the segmented images. Each cell was labeled with a unique identification number (Figure S3B). A vertex was determined by calculating the vertex degree, which gives information on the number of edges attached to one vertex. The number of vertices present in each cell is detected using pattern recognition. By going through the 4-connectivity binarized image, the sum of pixels within the 3×3 neighborhood of each foreground pixel (for a binary image, 1 is the foreground pixel and 0 is the background pixel) is determined. Therefore, the vertex degree *k* may be written as

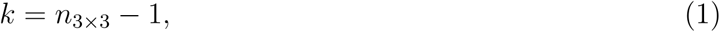

where *n*_3×3_ is the number of foreground pixels in the 3×3 neighborhood.

From a biological point of view, a vertex is where multiple (i.e. three or more) edges meet. Therefore, *k* has to be bigger or equal to 3 in order to be considered as a vertex point. This results in a polygonal lattice of cells, in which each of the polygons consists of a unique set of vertices and edges that are crucial for further cell shape and topology analysis as well as polarity quantification.

We then identified the immediate neighbors of individual segmented cells. This step serves as a pre-requisite for the measurement of neighbor vector polarity.

### Quantification of planar polarization using the Fourier Series method

Planar polarity quantification based on the Fourier Series method is implemented as described in (Aigouy et al., 2010; Merkel et al., 2014). Using the intensity of junctional proteins, the symmetric tensor components *Q*_1_ and *Q*_2_ are computed as described (see Equation 3–4 in (Aigouy et al., 2010)). In order to allow for polarity comparison between images, *Q*_1_ and *Q*_2_ are normalized to normalization constant *N* where 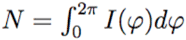 as follows:

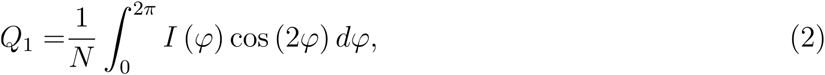

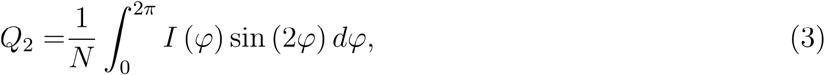

where *I*(*φ*) is the intensity of junctional proteins at segmented cell boundary at an angle *φ* defined with respect to the centroid of the cell.

The magnitude and angle of polarity in each cell were defined as described (Equation 5 in (Aigouy et al., 2010)).

In the Fourier Series method, polarity magnitude starts from 0 onwards, with 0 having complete zero polarization, while increasingly values represent increasing polarization. All polarity angles (in degree) range between −90° to +90°, with 0° corresponding to the x-axis of the image.

To allow direct comparison between all polarity methods, we computed the normalized Fourier Series polarity magnitudes by normalizing against its maximum value.

### Quantification of planar polarization using the Ratio method

Planar polarity quantification based on the Ratio method described in (Strutt et al., 2016) is implemented and improved as follows. Within each cell, junctional fluorescence intensity is grouped into four bins of equal size (90°). The binned data are then smoothed out using linear interpolation. The mean intensity that falls within the opposing pair of bins are summed up and the ratio between the bin pairs is denoted as asymmetry. All the asymmetries are rounded to a precision of 1e-3. Thus, the maximum asymmetry is the polarity magnitude. The central angle which corresponds to the average of all angles from multiple maximum asymmetries is considered as the polarity angle.

In the Ratio method, polarity magnitude starts from 1 onwards, with 1 having complete zero polarization, while increasingly value represents increasing polarization. All polarity angles (in degree) range between −90° to +90°, with 0° corresponding to the x-axis of the image.

The normalized Ratio polarity magnitude of a cell is computed by subtracting 1 from the ratio value and normalizing it against the maximum of all the subtracted ratio values from all cells. This allows for direct comparison between all polarity methods.

### Quantification of planar polarization using a PCA-based method

Here, we implemented a Principal Component Analysis-based method to quantify 2D planar polarization. First, fluorescence intensity *I*(*θ*) of the protein of interest is extracted from original image at an angle *θ* on the segmented cell boundary, with polar coordinates taken with respect to the centroid of the cell.

In order to mitigate the effect of denseness of points on the calculation of covariance matrix, the weighting *dθ_i_* for each *i* point on the cell boundary is calculated as follows:

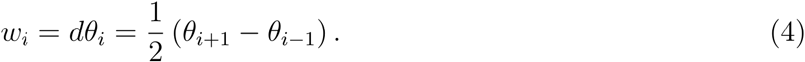

For each point *i* on the cell boundary with intensity *I_i_*, all the intensities are normalized so that it is independent of the image format (for example, 8-bit, 12-bit, 16-bit, and etc):

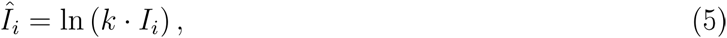

where *k* is the normalization factor (≈ 1000 empirically). The transformed coordinates, 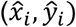, can then be determined as follow:

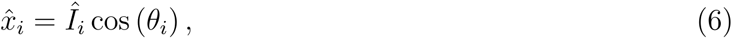

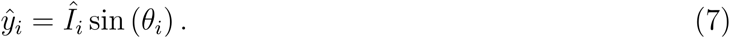

Next, the covariance matrix, ***σ***, is calculated as follows:

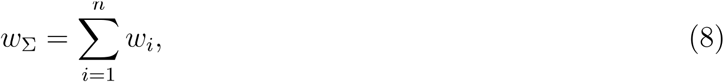

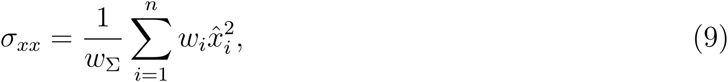

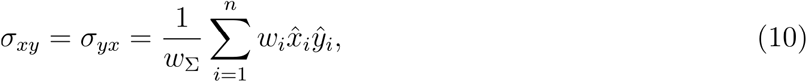

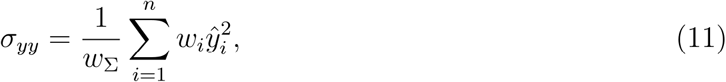

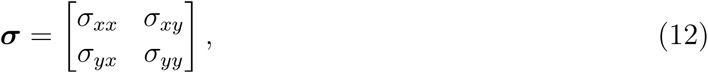

where *w*_Σ_ is the sum of all weightings and *σ_μv_* are the covariances.

Eigenvalues *λ*_1_, *λ*_2_ with *λ*_1_ ≥ *λ*_2_ and eigenvectors *v*_1_, *v*_2_ of the covariance matrix ***σ*** are computed accordingly. Using the eigenvalues and covariances, we defined the magnitude of polarity *p* and the angle of polarity *θ* for a single cell as

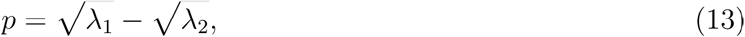

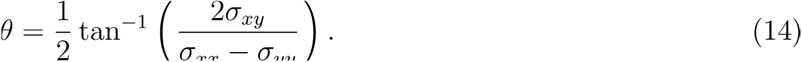

Polarity magnitude obtained from the PCA method starts from 0 onwards, with 0 having complete zero polarization, while increasingly values represent increasing polarization. The angle of polarity *θ* is measured with respect to the x-axis of an image. Polarity angle measurement ranges between −90° and +90°, with 0° oriented along the x-axis and ±90° oriented along the y-axis.

To allow direct comparison between all polarity methods, we computed the normalized PCA polarity magnitudes by normalizing against its maximum value.

### Quantification of planar polarization at tissue scales

The cell-by-cell polarity readout obtained from either method is further applied to measure local (neighbor and coarse-grain) polarization (Figure S3B’(ii-iii)). Within a group of cells, the polarity measurements can be combined in specific ways to reveal the strength of polarity as well as the polarity coordination between cells. The most direct way is to compute the average of polarity magnitude without taking its polarity angle into consideration, which is termed the “Average Polarity Magnitude”. This measure gives us an idea of the average polarization strength for all cells within the image. It can be simply computed with the following equation:

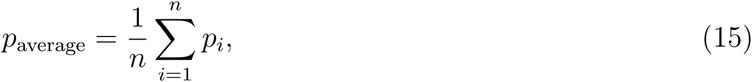

where *p_i_* is the polarity magnitude of *i*-th cell and *n* is the total number of cells.

For the described simulation and experimental data, we used the Average Polarity Magnitude measure as a simple readout of polarity strength.

On the other hand, “vector” average polarity measurement is defined to capture the strength and coordination of planar polarity between groups of cells within a defined area (referred to as “Coarse-Grain Vector Polarity”) and with its immediate neighbors (referred to as “Neighbor Vector Polarity”). First, polarity of individual cells is converted into their vector form 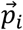. The vector polarity average, 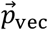, is computed as follows:

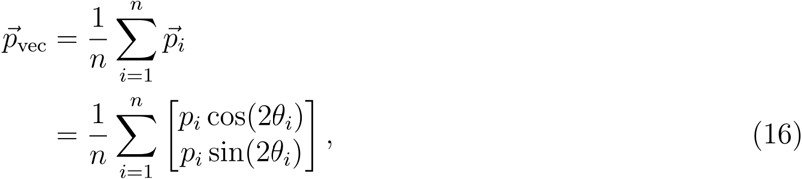

where 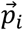, *p_i_*, *θ_i_* are the polarity vector, polarity magnitude and polarity angle of *i*-th cell respectively. *n* is the total number of cells. Therefore, vector average polarity magnitude *p*_vec_ and angle *θ*_vec_ can be explicitly written as

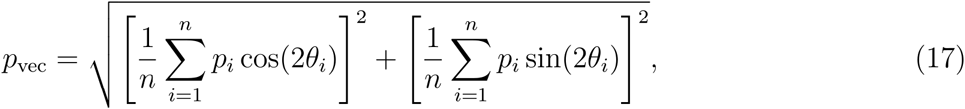

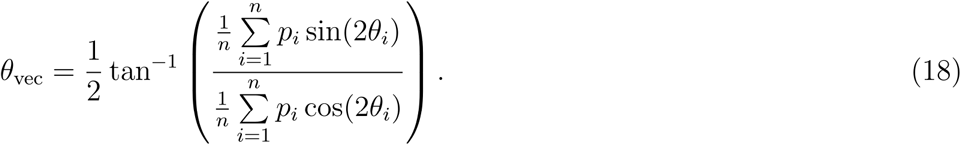

Notice that the computed vector average polarity magnitude *p*_vec_ takes the polarity angle of individual cells into consideration. Therefore, *p*_vec_ ≤ *p*_average_ for all possible cases of polarity magnitudes and angles, with equality if and only if all polarity angles are equal.

To visualize the coarse-grain polarity on the scale of groups of cells or the entire image area, cells are divided into groups with equal number of cells. In each group of cells, the vector average magnitude *p*_vec_ and angle of polarity *θ*_vec_ are computed as described in Equation 17 and 18 (Figure S3B’(ii)). On the other hand, to capture local polarity coordination of individual cells with its immediate neighbors, each cell’s immediate neighbors are identified and computed for vector average magnitude *p*_vec_ and angle of polarity *θ*_vec_ as described in Equation 17 and 18 (Figure S3B’(iii)). The neighbor vector polarity magnitudes obtained from all the cells are averaged across the tissue to obtain the average neighbor vector polarity magnitude measure.

Apart from that, circular statistics are used to quantify the degree of alignment or coordination of polarity angle between cells. A measure called circular angle variance as implemented in CircStats MATLAB toolbox (Berens, 2009) is used to determine the circular spread of vectorial data. In order to accommodate the rotational symmetry of polarity angle, the angle variance Var_circ_ for polarity angles can be computed as follows:

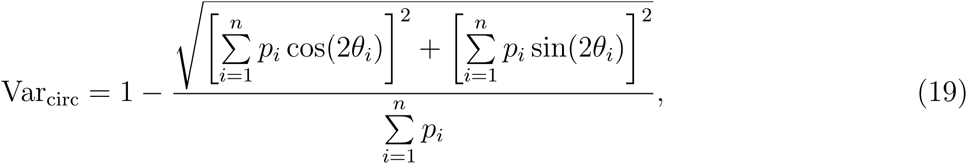

where *θ*_*i*_ is the polarity angle of *i*-th cell and *n* is the total number of cells.

Angle variance ranges between 0 and 1, with 0 having complete agreement in polarity alignment, while 1 represents complete polarity misalignment.

Additionally, we computed the mean angle difference, 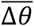, as a way to quantitatively compare the polarity angles obtained from the PCA against the Ratio and Fourier Series methods. Briefly, this is done first by calculating the angle differences between two polarity methods, accommodating the circular spread of the data. We then computed the mean of the (absolute) angle difference, with 0° representing complete agreement between polarity angles obtained by two different methods, while higher values indicate that the methods agree less.

### Circular weighted histogram

To visually compare cell polarity angle obtained from different planar polarity methods, we plotted a circular weighted histogram described as follow. First, we computed the magnitude and angle/axis of polarity on a cell-by-cell basis using all three methods. Data from multiple wings were combined and represented by a circular weighted histogram using the MATLAB built-in function *“polarhistogram"*. Data was grouped into 20 bins, with each bin representing a unique polarity angle. Histograms were weighted by the average magnitude of polarity within each bin to capture both the angle and magnitude of polarity (Aw et al., 2016). The length of each bin represents the polarity magnitude-weighted frequency of occurrences, meanwhile the orientation of each bin represents the axis of average polarity (results obtained from Ratio, Fourier Series and PCA methods were labeled in magenta, green and blue respectively). Note that the angle of polarity has characteristic rotational symmetry, hence a polarity angle of *θ* also corresponds to *θ* + *π* .nTherefore, for better visual representation, all computed polarity angles (ranges from −90° and +90°) were plotted in a range between 0° to 360°, with 0° corresponds to the x-axis of the image.

### Cell morphological parameter measurements

#### Cell area and perimeter

The apical cell area (pixel^2^) and perimeter (pixel) for each cell was determined from the labeled images using the MATLAB built-in function “*regionprops*”.

#### Cell shape regularity

Cell shape regularity was quantified based on how “far” a cell is from a regular polygon, using a measure focusing on the equilateral and equiangular properties of a polygon (Chalmeta et al., 2013). From the lengths of the edges (*l*_*i*_), first the median length of edges (*l*_*median*_) and the sum of all edge lengths (*l*_Σ_) was determined. Then, an intermediate term *D* can be calculated as follows:

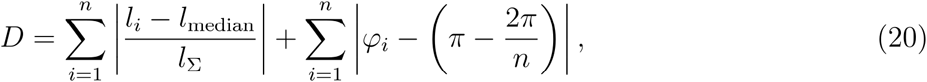

where *n* is the number of sides, and *φ_i_* are the interior angles of a cell. Finally, cell shape regularity measure *μ* can be obtained as follow:

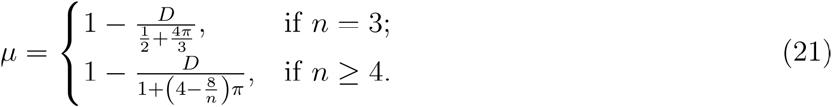

The value of cell regularity (a.u.) ranges from 0 to 1, with 0 represent highly irregular and 1 being perfectly regular with equal length of cell edges and interior angles.

#### Cell eccentricity and orientation

To measure cell eccentricity, a robust ellipse fitting approach was used. It is a shape-based method where the cell boundaries are used as a reference landmark for ellipse fitting (Young, 2010). Any ellipse can be described by the following (general) equation:

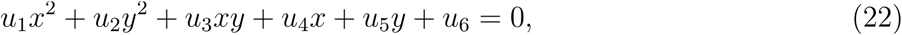

where *u_i_* is the unique coefficients of each distinct ellipse and (*x*, *y*) are the coordinates of the cell boundaries. The least squares method is used to determine the most optimal set of coefficients *u_i_* for every single cell. Then, the parameters of an ellipse can be determined using the following equations:

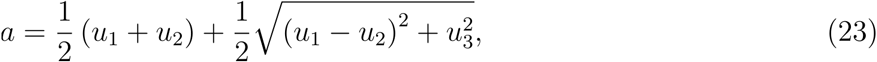

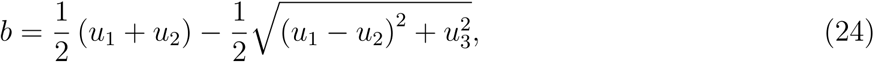

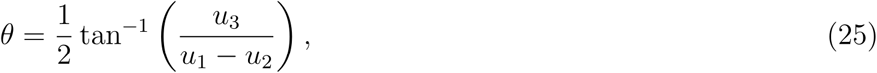

where *a*, *b* are the semi-major and semi-minor axes respectively with *a* ≥ *b*, and *θ* represents the cell orientation. The value for cell orientation ranges from −90° to +90°, with 0° corresponding to the x-axis of the image.

By fitting an ellipse onto the geometry of a cell, the eccentricity can be calculated using the following formula:

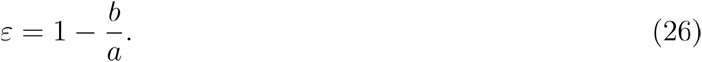

The value for cell eccentricity *ε* ranges from 0 to 1 in arbitrary units, with 0 represent no elongation (or circular) and 1 being highly eccentric.

#### Number of cell junctions

A cell junction that is below 10% of average junctional length will not be considered as a side of a cell. Once cell junctions are defined, the number of cell junctions which is equivalent to number of neighboring cells can be determined.

## Acknowledgements

We thank Natalia Bulgakova and Lindsay Canham for kindly providing *Drosophila* embryonic epidermal and wing disc images and Samantha Warrington and Aydar Uatay for comments on the manuscript. We thank Suzanne Eaton and the Bloomington Drosophila Stock Center for fly stocks. Excellent technical support was provided by the Department of Biomedical Science Fly Facility and imaging was performed in the Wolfson Light Microscopy Facility.

## Author Contributions

ST and WT developed the PCA method. KF implemented the Fourier Series method. DS developed the Ratio method. ST designed and developed the QuantifyPolarity GUI and simulations, improved the Ratio method and conducted all experiments and analysis. ST and DS wrote the manuscript. DS conceived the project.

## Data Availability

An installer for the QuantifyPolarity GUI for Mac and PC, a user manual and links to video tutorials can be downloaded from https://github.com/QuantifyPolarity/QuantifyPolarity.

---

**Figure S1.**
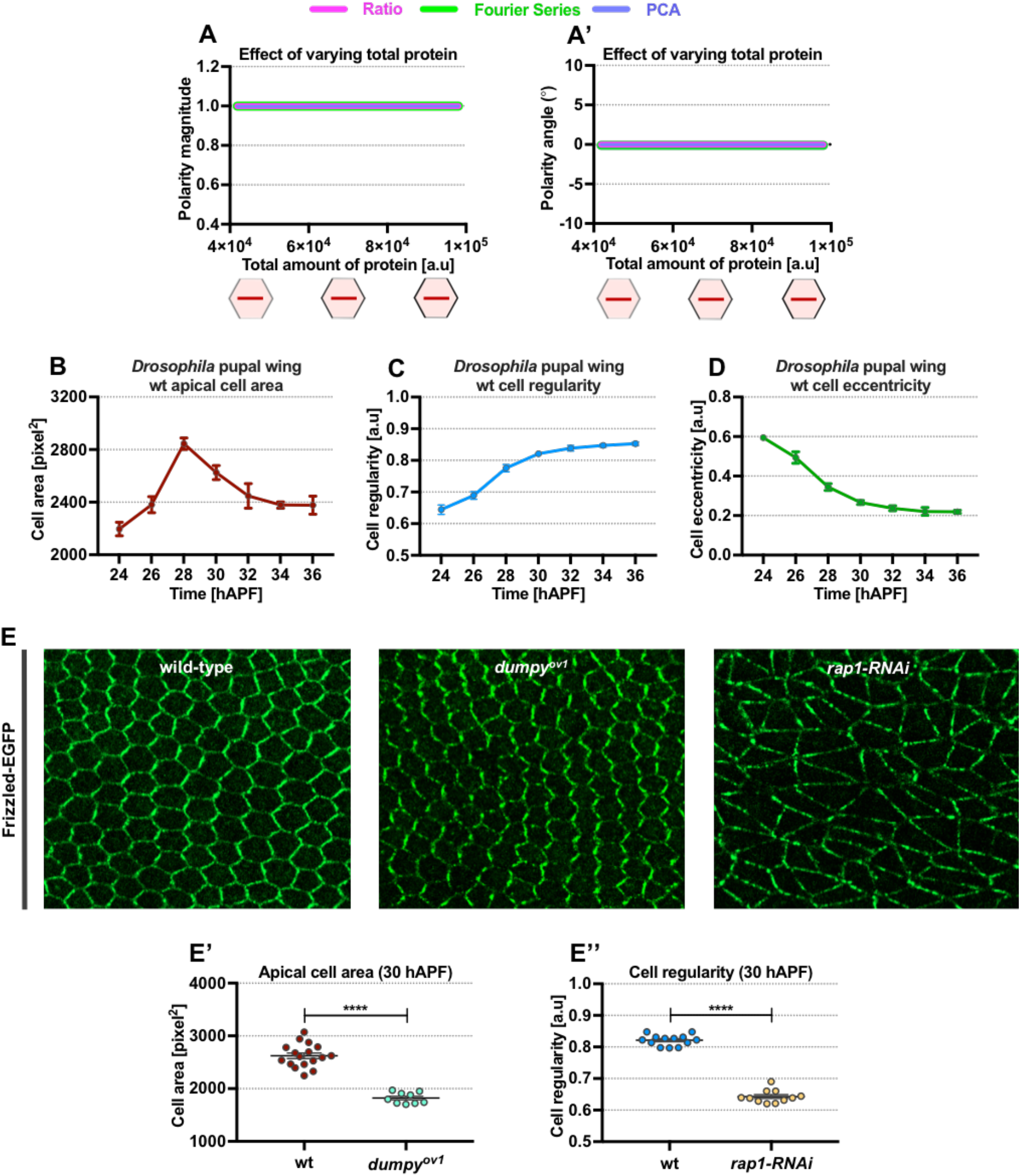
related to Figure 1. The variation of cell size and shape during pupal wing development and in different genotypes. (A-A’) Effects of varying total amount of proteins on the entire cell junctions with conserved cell geometries and junctional protein distribution. Graphs show quantified polarity magnitudes (A) and polarity angles (A’) obtained from varying the amount of proteins on cell junctions. All polarity magnitudes (a.u.) obtained using each different method are normalized. All polarity angles (in degree) range between −90° to +90°, with 0° corresponding to the x-axis of the image. (B-D) Quantification of average (C) apical cell area, (D) cell regularity, (E) cell eccentricity of otherwise wild-type pupal wings expressing Fz-EGFP from 24 to 36 hAPF (*n* = 4 - 5 wings analyzed). (E) Confocal images of otherwise wild-type, *dumpy^ov1^* mutant and *rap1-RNAi* pupal wings expressing Fz-EGFP at 30 hAPF. *dumpy^ov1^* mutant wings lack the extracellular matrix protein Dumpy required to regulate proper wing shape and size development. *rap1-RNAi* wings lack the homogeneous distribution of E-Cadherin required for regulation of cell shape. Note distinctive sizes and shapes of planar polarized cells on these mutant backgrounds as compared to otherwise wild-type cells. (E’) Quantified average apical cell area of *dumpy^ov1^* and wild-type wings at 30 hAPF (*n* = 11 - 13 wings per genotype analyzed). (E’’) Quantified average cell regularity of *rap1-RNAi* and wild-type wings at 30 hAPF (*n* = 11 - 13 wings per genotype analyzed). Error bars indicate mean±SEM. Unpaired t-test. Significance levels: p-value ≤ 0.0001^****^ and p-value ≤ 0.0067^**^.

**Figure S2.**
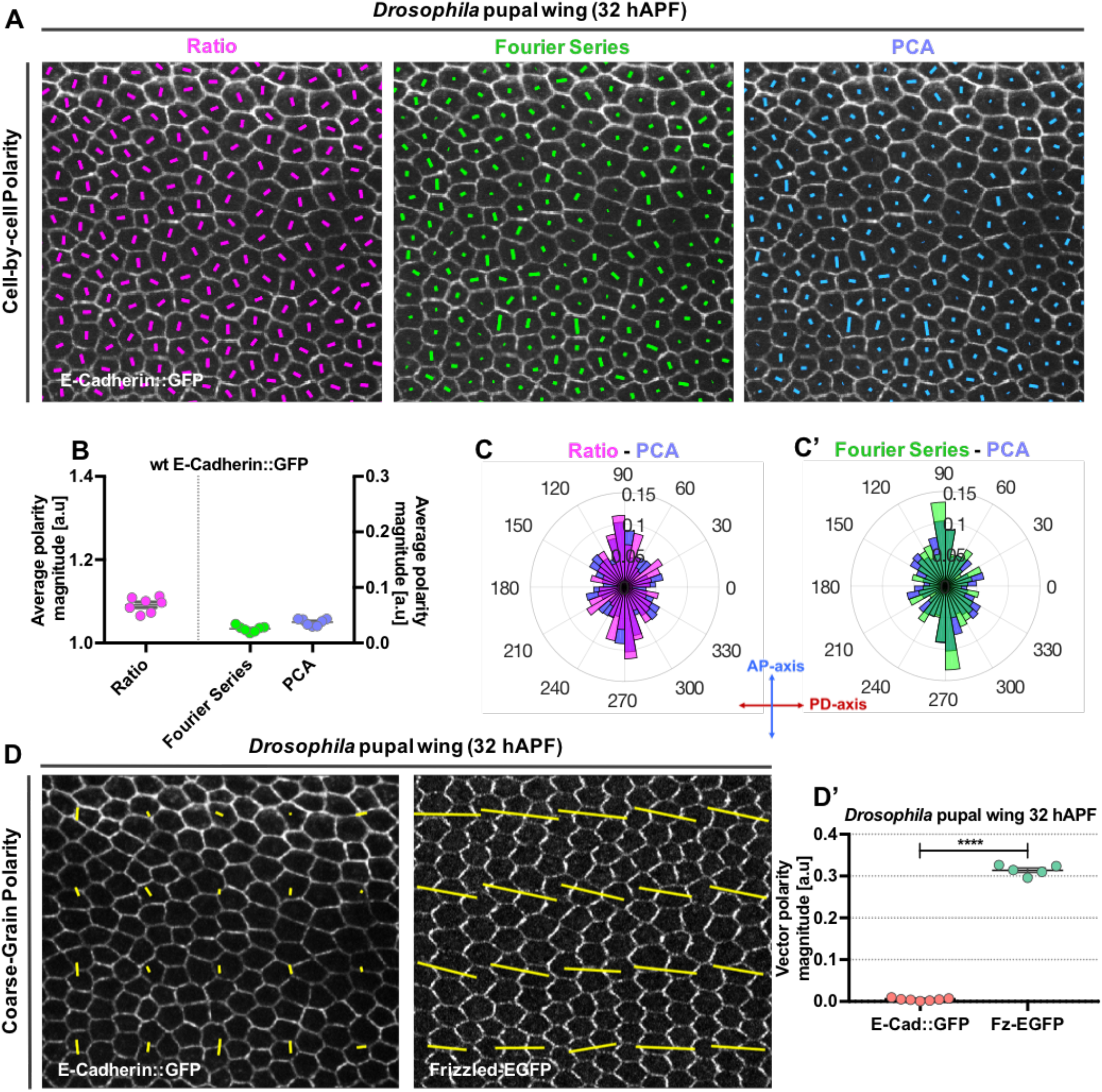
related to Figure 2. The weakly polarized distribution of E-Cadherin on cell junctions results in low polarity magnitude with dispersed polarity angles. (A) Quantified cell-scale polarity pattern of otherwise wild-type wings expressing E-Cadherin::GFP at 32 hAPF using three different methods. The blue, green, magenta bars represent the magnitude (length of bar) and angle (orientation of bar) of planar polarization pattern for a given cell obtained from PCA, Fourier Series and Ratio methods respectively. (B) Plot of average polarity magnitudes at 32 hAPF obtained from PCA, Fourier Series and Ratio methods respectively (*n* = 7 wings analyzed). Average polarity magnitudes with its standard deviation obtained using each different method are indicated. Error bars indicate mean±SEM. (C-C’) Circular weighted histogram plots displaying the orientation of E-Cadherin::GFP polarity obtained from (C) Ratio and PCA and (C’) Fourier Series and PCA (*n* = 7 wings analyzed). (D) Quantified coarse-grain polarity pattern of otherwise wild-type wings expressing E-Cadherin::GFP and Frizzled-EGFP at 32 hAPF. The yellow bars represent the magnitude (length of bar) and angle (orientation of bar) of planar polarization pattern for a group of cells obtained from the PCA method. (D’) Plot of vector average polarity magnitude for E-Cadherin::GFP and Frizzled-EGFP expressing wild-type wings at 32 hAPF (*n* = 5 to 7 wings analyzed). Error bars indicate mean±SEM. Unpaired t-test. Significance levels: p-value ≤ 0.0001^****^.

**Figure S3.**
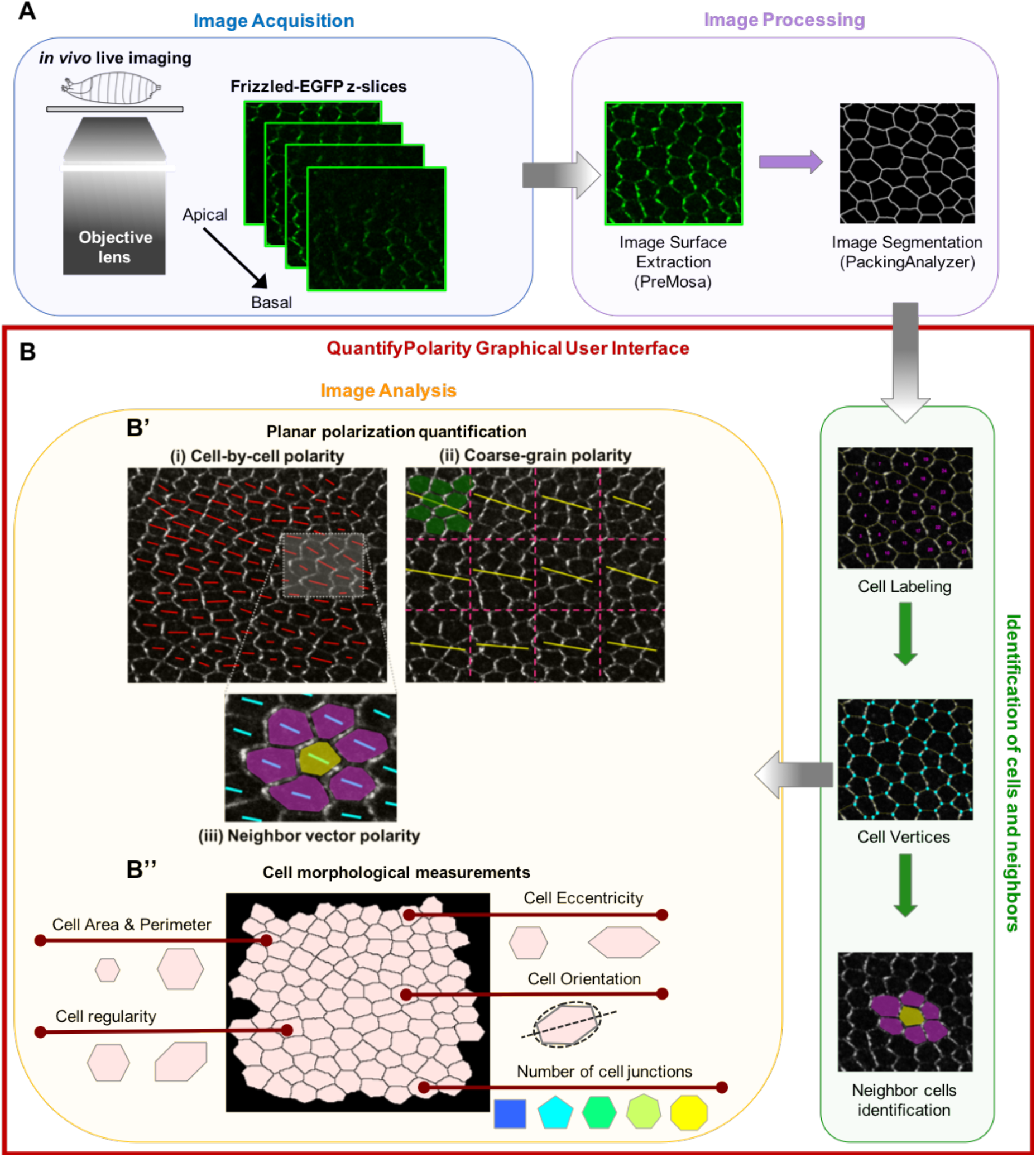
related Figure 4. Overview of image acquisition, processing and subsequent image analysis steps. (A) Following image acquisition, raw images are processed by external tools (e.g. PreMosa and PackingAnalyzer) to obtain segmented images. These images are then fed into QuantifyPolarity GUI. (B) Following identification of cells and their neighbor relations (Green box), QuantifyPolarity performs further image analysis (Yellow Box), such as (1) polarity and (2) cell morphology quantifications. (B’) Quantification of planar polarity at cellular and tissue scales. (i) Cell-by-cell polarity pattern of a *Drosophila* pupal wing expressing Fz-EGFP at 30 hAPF. The length and orientation of red bars denote the polarity magnitude and angle for a given cell respectively. (ii) Coarse-grain pattern of vector average polarity at 30 hAPF. Image is divided into group of cells with equal square grids (with dotted magenta lines), where the vector average polarity for each group of cells is computed. For each group of cells, the average polarity magnitude *p*_vec_ is proportional to the length of the yellow bar, while the average polarity angle *θ*_vec_ is denoted by the orientation of the yellow bar. (iii) Neighbor vector polarity quantification for yellow cell with its immediate neighbors (magenta cells). The length and orientation of cyan bars denote the polarity magnitude and angle for yellow cell. (B’’) Cell morphological measurement tools are available in QuantifyPolarity GUI to quantify cell area, perimeter, shape regularity, eccentricity, orientation and number of cell junctions.

**Figure S4.**
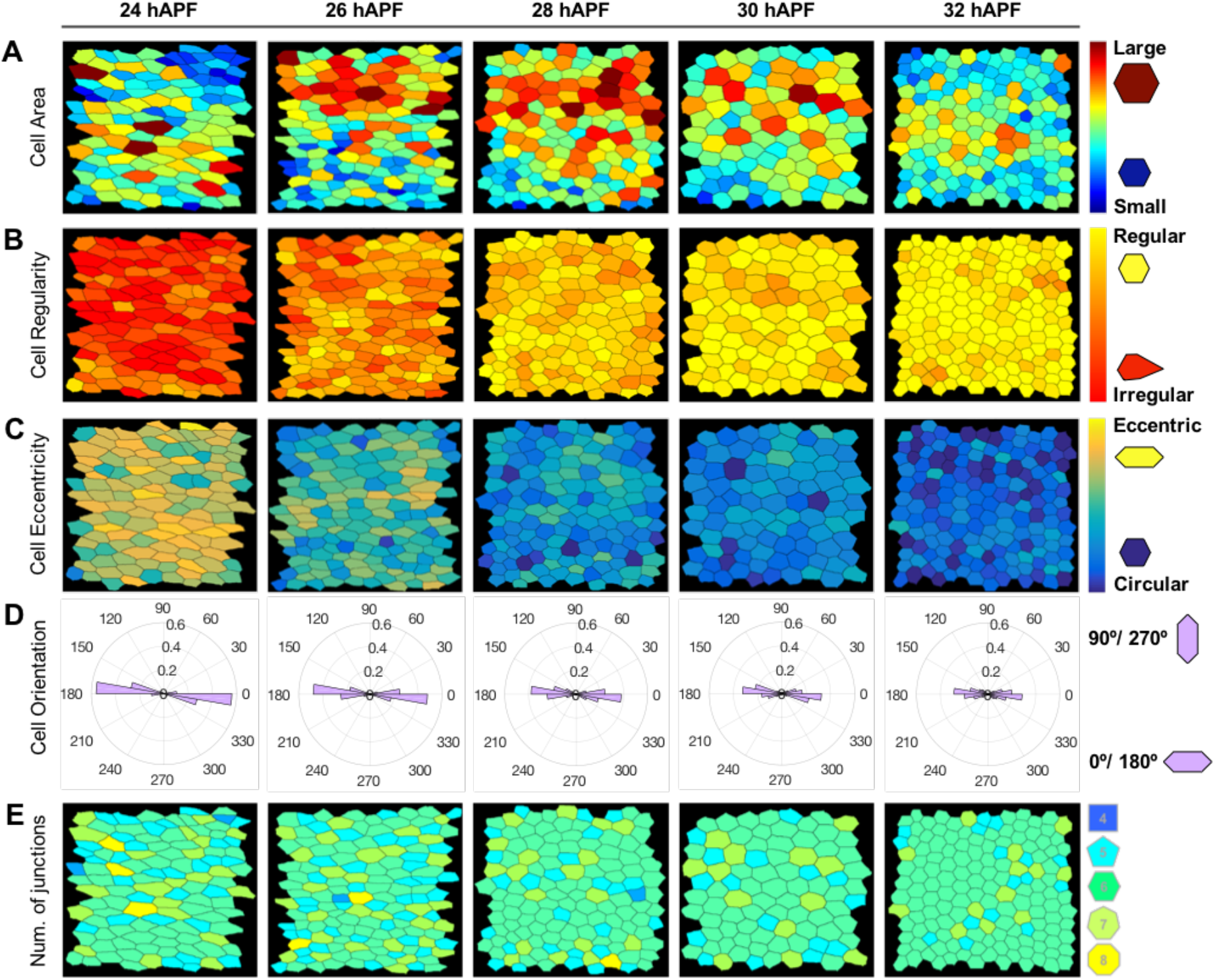
related to Figure 4. Examples of cell morphological quantitative analysis of *Drosophila* pupal wing development computed using QuantifyPolarity GUI. (A-E) Processed images of the posterior-proximal region of otherwise wild-type pupal wing from 24 to 32 hAPF. (A) Cells are color-coded according to the cell apical area, with red represent cells with larger apical area and blue represents cells with smaller apical area. (B) Cells are color-coded according to the regularity of the shape, with yellow being perfectly regular and red represent highly irregular. (C) Cells are color-coded according to the eccentricity of the shape, with yellow represent highly eccentric and blue being circular or non-eccentric. (D) Circular histogram plots display the orientation of cell, ranges between 0° to 360°, with 0°/180° corresponding to the x-axis of the image. (E) Cells are color-coded according to the number of cell junctions.

**Supplementary Table 1.**
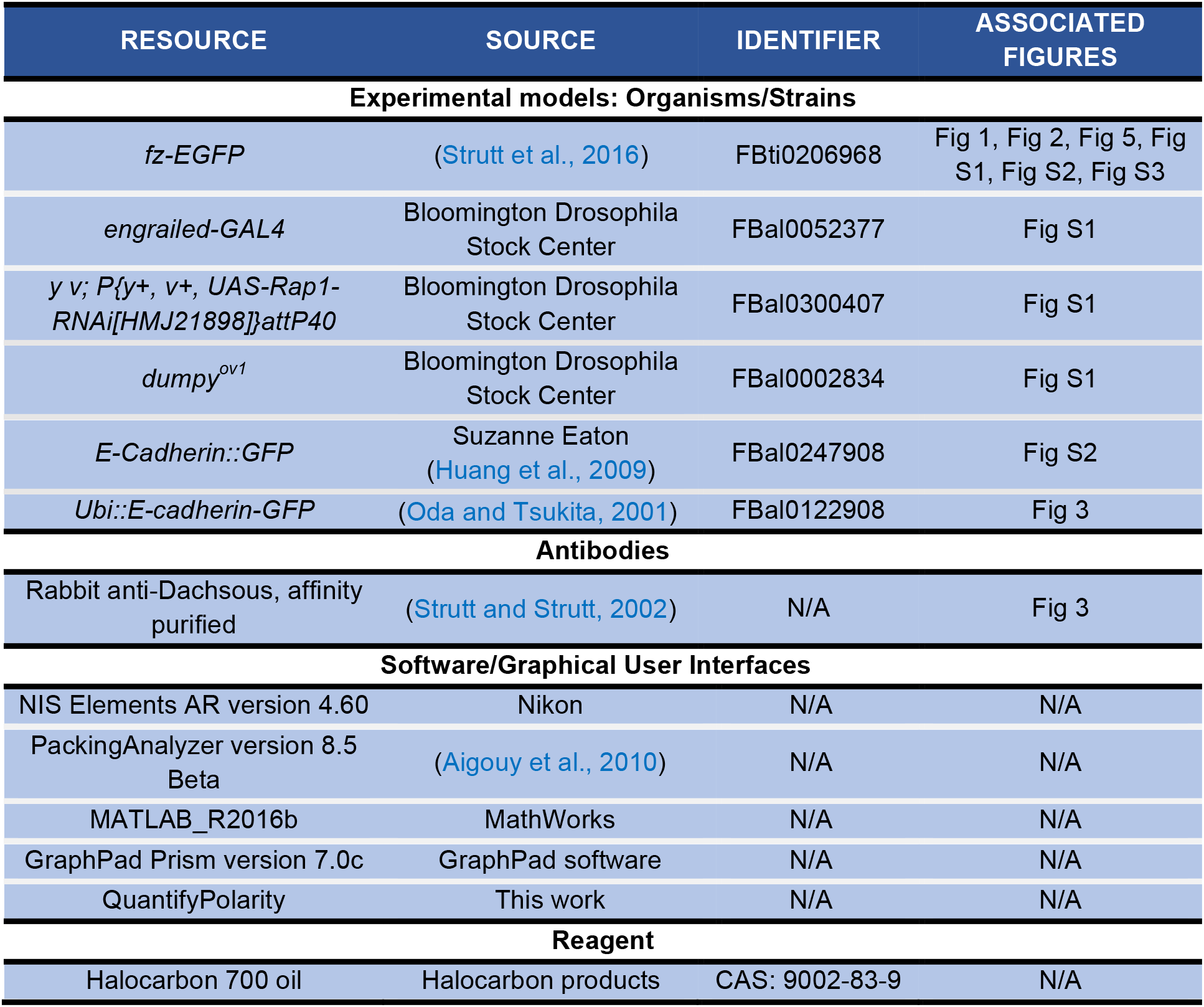
Key resources

| RESOURCE | SOURCE | IDENTIFIER | ASSOCIATED FIGURES |
| --- | --- | --- | --- |
| **Experimental models: Organisms/Strains** |  |  |  |
| *fz-EGFP* | (Strutt et al., 2016) | FBti0206968 | Fig 1, Fig 2, Fig 5, Fig S1, Fig S2, Fig S3 |
| *engrailed-GAL4* | Bloomington Drosophila Stock Center | FBal0052377 | Fig S1 |
| *y v; P{y+, v+, UAS-Rap1-RNAi[HMJ21898]}attP40* | Bloomington Drosophila Stock Center | FBal0300407 | Fig S1 |
| *dumpy<sup>ov1</sup>* | Bloomington Drosophila Stock Center | FBal0002834 | Fig S1 |
| *E-Cadherin::GFP* | Suzanne Eaton<br>(Huang et al., 2009) | FBal0247908 | Fig S2 |
| *Ubi::E-cadherin-GFP* | (Oda and Tsukita, 2001) | FBal0122908 | Fig 3 |
| **Antibodies** |  |  |  |
| Rabbit anti-Dachsous, affinity purified | (Strutt and Strutt, 2002) | N/A | Fig 3 |
| **Software/Graphical User Interfaces** |  |  |  |
| NIS Elements AR version 4.60 | Nikon | N/A | N/A |
| PackingAnalyzer version 8.5 Beta | (Aigouy et al., 2010) | N/A | N/A |
| MATLAB_R2016b | MathWorks | N/A | N/A |
| GraphPad Prism version 7.0c | GraphPad software | N/A | N/A |
| QuantifyPolarity | This work | N/A | N/A |
| **Reagent** |  |  |  |
| Halocarbon 700 oil | Halocarbon products | CAS: 9002-83-9 | N/A |

